# Co-translational folding allows misfolding-prone proteins to circumvent deep kinetic traps

**DOI:** 10.1101/721613

**Authors:** Amir Bitran, William M. Jacobs, Xiadi Zhai, Eugene Shakhnovich

## Abstract

Many large proteins suffer from slow or inefficient folding *in vitro*. Here, we provide evidence that this problem can be alleviated *in vivo* if proteins start folding co-translationally. Using an all-atom simulation-based algorithm, we compute the folding properties of various large protein domains as a function of nascent chain length, and find that for certain proteins, there exists a narrow window of lengths that confers both thermodynamic stability and fast folding kinetics. Beyond these lengths, folding is drastically slowed by non-native interactions involving C-terminal residues. Thus, co-translational folding is predicted to be beneficial because it allows proteins to take advantage of this optimal window of lengths and thus avoid kinetic traps. Interestingly, many of these proteins’ sequences contain conserved rare codons that may slow down synthesis at this optimal window, suggesting that synthesis rates may be evolutionarily tuned to optimize folding. Using kinetic modelling, we show that under certain conditions, such a slowdown indeed improves co-translational folding efficiency by giving these nascent chains more time to fold. In contrast, other proteins are predicted not to benefit from co-translational folding due to a lack of significant non-native interactions, and indeed these proteins’ sequences lack conserved C-terminal rare codons. Together, these results shed light on the factors that promote proper protein folding in the cell, and how biomolecular self-assembly may be optimized evolutionarily.

**Significance Statement:** Many proteins must adopt a specific structure in order to perform their functions, and failure to do so has been linked to disease. Although small proteins often fold rapidly and spontaneously to their native conformations, larger proteins are less likely to fold correctly due to the myriad incorrect arrangements they can adopt. Here, we show that this problem can be alleviated if proteins start folding while they are being translated, namely, built one amino acid at a time on the ribosome. This process of co-translational folding biases certain proteins away from misfolded states that tend to hinder spontaneous refolding. Signatures of unusually slow translation suggest that some of these proteins have evolved to fold co-translationally.

---

Many large proteins refold from a denatured state very slowly *in vitro* (on timescales of minutes or slower) while others do not spontaneously refold at all (1–6). Given that proteins must rapidly and efficiently fold in the crowded cellular environment, how is this conundrum resolved? The answer likely involves a number of factors that affect cellular folding, but which are absent in vitro. For example, molecular chaperones such as GroEL in *E. Coli*, and TriC and HSP90 in eukaryotes may substantially improve folding efficiency by confining unfolded chains to promote their folding, or by repeatedly binding and unfolding misfolded chains until the correct strucure is attained (6–11). A second, more recently appreciated factor that may improve *in vivo* folding efficiency is co-translational folding on the ribosome (12–19), which may affect the folding of as much as 30% of the E. Coli proteome (19). A recent set of works (12, 13) suggests that protein synthesis rates in various organisms may be under evolutionary selection to allow for co-translational folding. Namely, these works show that conserved stretches of rare codons, which are typically translated more slowly than their synonymous counterparts, are significantly enriched roughly 30 amino acids upstream of chain lengths at which folding is predicted to begin. This 30 amino acid gap is expected given that the ribosome exit tunnel sequesters the last ∼ 30 amino acids of a nascent chain and generally impedes their folding. The observed correlation between chain lengths that allow for folding and conserved rare codons suggests that co-translational folding may be under positive evolutionary selection. However, the specific mechanisms by which co-translational folding is beneficial have not been elucidated.

Here, we address this question using an all-atom computational method for inferring detailed protein folding pathways and rates while accounting for the possibility of non-native conformations. We apply this method to compute folding properties of proteins at various nascent chain lengths to address how the vectorial nature of protein synthesis may affect co-translational folding efficiency. We find that for certain large proteins, vectorial synthesis is beneficial because it allows nascent chains to fold rapidly at shorter chain lengths, prior to the synthesis of C-terminal residues which stabilize non-native kinetic traps. Many of these proteins’ sequences contain conserved rare codons ∼30 amino acids downstream of these faster-folding intermediate lengths, suggesting these protein sequences may have evolved to provide enough time for co-translational folding. We also identify counterexamples—proteins without conserved rare codons that do not misfold into deep kinetic traps, and for which vectorial synthesis thus confers no advantage. Together, these results shed light on how biophysical folding properties of nascent chains determine the advantages of co-translational folding, and how co-translational folding may be optimized evolutionarily.

## Results

### Predicting folding properties of nascent chains

In order to compute co-translational folding pathways and rates, we developed a simulation-based method and analysis pipeline described in Fig. 1 and Methods. The method utilizes an all-atom Monte-Carlo simulation program with a knowledge-based potential and a realistic move-set described previously (20–22). In essence, rather than simulating a protein’s folding ab initio from an unfolded ensemble (which is intractable for large proteins at reasonable simulation timescales), we simulate *unfolding*, and in tandem, calculate the free energies of the folded, unfolded and various intermediate states from simulations with enhanced sampling. Given rates of sequential unfolding between these states and their free energies, the reverse folding rates can be computed from detailed balance. Importantly, our sequence-based potential energy function is not biased towards the native state, as in native-centered (Gō) models, and allows for the possibility of non-native interactions. Thus we can account for the role of misfolded states in folding kinetics. This method is applied at multiple chain lengths to predict co-translational folding properties.

**Fig. 1.**
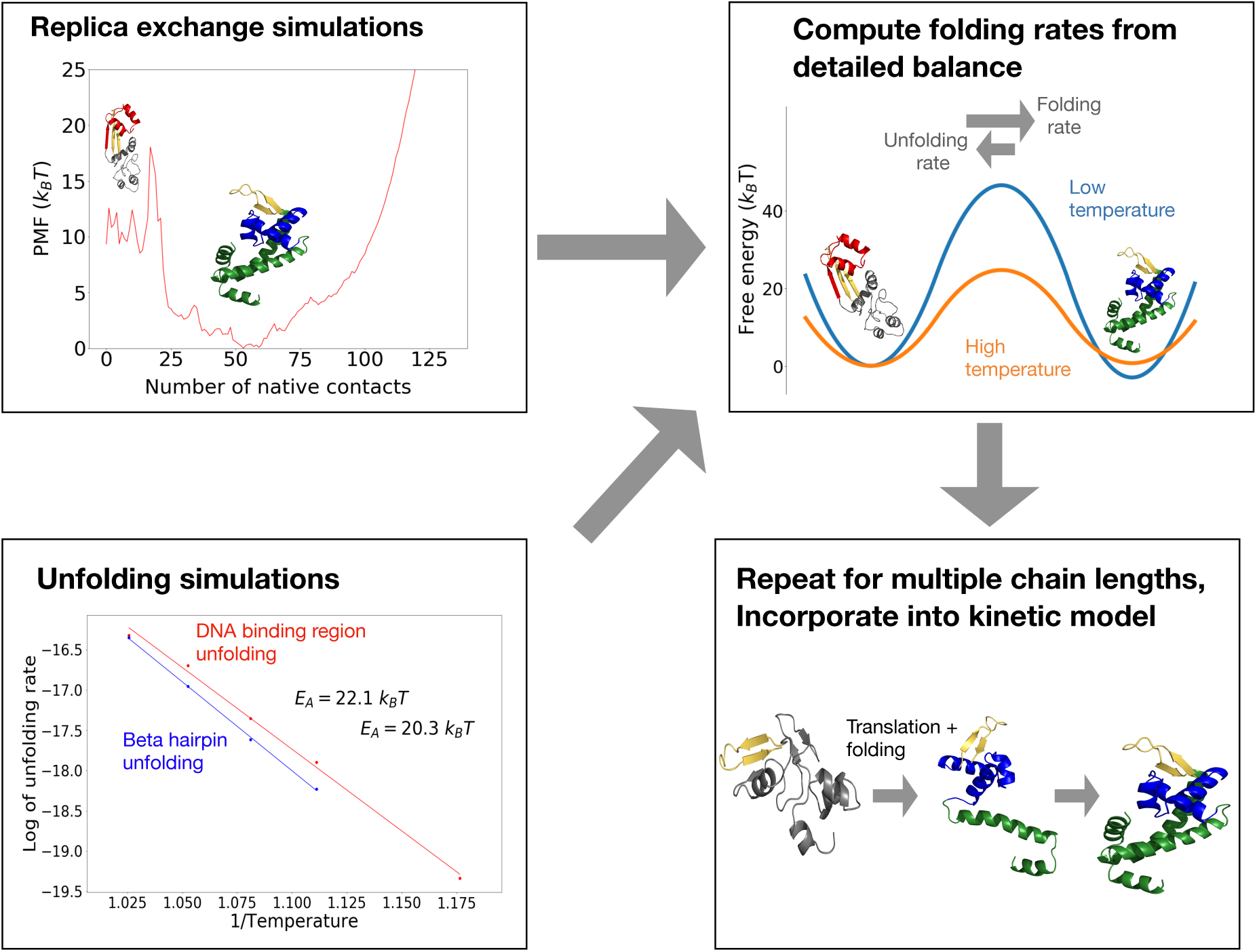
(Top let) We run replica exchange atomistic simulations with a knowledge-based potential and umbrella sampling to compute a protein’s free energy landscape. (Bottom left) To obtain barrier heights, we run high-temperature unfolding simulations and extrapolate unfolding rates down to lower temperatures assuming Arrhenius kinetics. (Top right) The principle of detailed balance is then used to compute folding rates. (Bottom left) The process is repeated at multiple chain lengths and incorporated into a kinetic model of co-translational folding. For details, see Methods.

Our approach here is based on a few key assumptions: 1.) The ribosome will not significantly affect co-translational folding pathways, and thus is neglected. Previous work suggests that the ribosome’s destabilizing effect on nascent chains is relatively modest, typically 1-2 kcal/mol (23), and affects various folding intermediates to a comparable extent (24). Thus, the ribosome is expected not to drastically affect the relative stability of the different intermediates computed here. 2.) Unfolding rates are assumed to obey Arrhenius kinetics, such that rates computed at high temperatures can be readily extrapolated to lower temperatures. This is justifiable so long as the barriers between intermediates are large so that a local equilibrium is reached in each free energy basin prior to unfolding. 3.) We assume that non-native contacts form on timescales faster than the timescales of native folding transitions. This assumption implies that a protein’s folding landscape can be described by macrostates characterized by certain folded native elements in fast equilibrium with non-native contacts that are compatible with the currently folded elements, and that these macrostates obey detailed balance (see Methods). This assumption holds in general for the misfolded states observed here, which are dominated by short-range interactions that form rapidly compared to the long-range contacts that stabilize most native structures.

### MarR-an *E. coli* protein with conserved rare codons-adopts stable co-translational folding intermediates

We began by simulating the co-translational folding of a protein previously shown to contain a conserved rare codons 30 amino acids downstream of a possible co-translational folding intermediate (12): the *E. Coli* Multiple Antibiotic Resistance Regulator (MarR). MarR, a transcriptional repressor (25–27), natively assembles into a winged helix homodimer with each monomer composed of a DNA binding region and a helical dimerization region (Fig. 2A). To investigate whether individual monomers are stable, we ran equilibrium replica exchange simulations with umbrella sampling using our all-atom potential (Methods). We find that the dimerization region is folded a fraction of the time, while the DNA binding region is stably folded the majority of the time at temperatures below *T* ≈ 0.9 *T*_*M*_ (blue dotted line), where *T*_*M*_ is the monomer melting temperature (see also Fig. S1B). These results indicate that the monomer acquires a substantial amount of native structure in isolation.

**Fig. 2.**
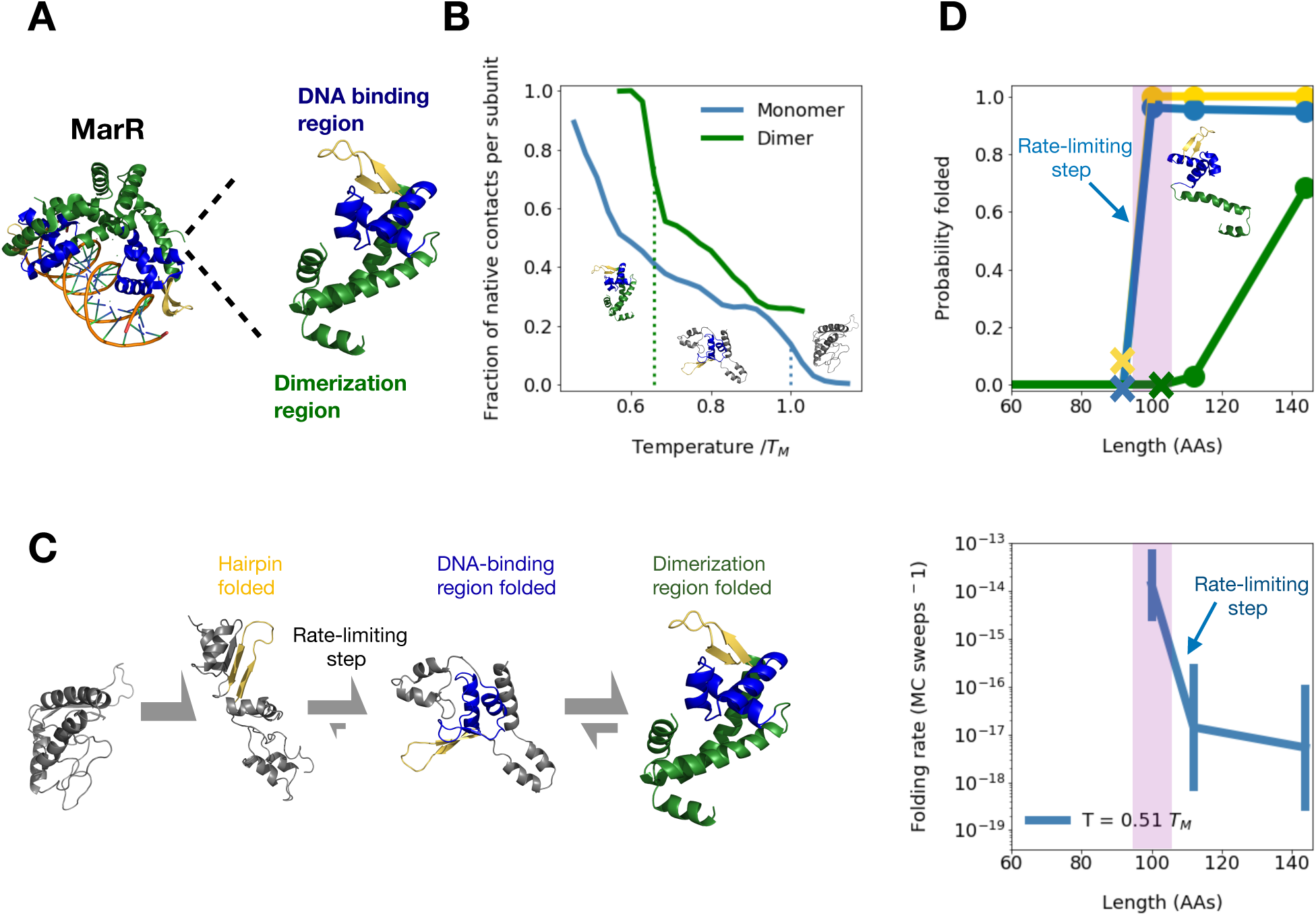
(A) Structure of native MarR dimer bound to DNA (left) as well as monomer (right) with highlighted dimerization region (green), DNA binding region (blue), and a crucial beta hairpin involved in stabilizing the DNA binding region (gold). (B) Mean fraction of native contacts per subunit for monomeric and dimeric MarR as a function of temperature normalized by DNA binding region melting temperature (right dashed line). The dimer melting temperature is indicated by the left dashed line. Sample monomeric structures from each temperature range are shown, illustrating melting of the dimerization region followed by the DNA binding region (C) Predicted folding pathway of MarR monomer. (See text for details.) (D) (Top) At various chain lengths, we plot the equilibrium probability that the structural elements associated with each folding step in the MarR monomer folding pathway are folded (gold = hairpin folding, blue = DNA binding region folding, green = dimerization region folding). X’s indicate the minimum chain lengths at which each step is possible. (Bottom) For each chain length shown in the top panel, we plot the rate of the slowest folding step–DNA-binding region formation. A narrow window of chain lengths that confers both folding speed and stability is highlighted in purple. Error bars on folding rates are obtained from bootstrapping. (see Methods) Both panels are shown at a simulation temperature of *T* = 0.51*T*_*M*_

We next turned to investigating the monomer’s folding pathway. We find that the monomer folds in three steps (Fig. 2C) characterized by: 1.) the relatively fast folding of a crucial beta hairpin composed of residues valine 84 through leucine 100 (gold in Fig. 2), which scaffolds the entire DNA binding region in the final structure, 2.) The completion of DNA binding region folding, which is the rate-limiting step involving the formation of long range contacts between one of the strands in the beta hairpin–leucine 97 through leucine 100– and another strand composed of alanine 53 through threonine 56 (blue in Fig. 2), and finally 3.) Folding of the dimerization region (green in Fig. 2), which is reversible as the helices comprising this region rapidly exchange between various native and non-native tertiary arrangements (Fig. S1B). Naturally, the dimerization region becomes substantially more ordered in the presence of a dimeric partner. Rates for each folding step as a function of temperature are shown in Fig. S2.

Having predicted the monomer’s folding pathway, we wondered whether these folding steps can take place cotranslationally. To test this, we truncated residues from the C-terminus of the protein and ran equilibrium simulations of the resulting nascent-like chains at various lengths. At each length, we computed the probability that the tertiary contacts associated with each folding step are formed at equilibrium (Fig. 2D top panel, see Methods for details), We find that as soon as the crucial beta hairpin (gold in Fig. 2) has been fully synthesized at length 100, both beta-hairpin folding and the rate-limiting DNA binding region folding step become thermodynamically favorable, suggesting folding can begin co-translationally at this length (see also Fig. S1F). This finding is in agreement with prior analysis using a coarse-grained model, which predicts a co-translational folding intermediate at a similar chain length (Fig. S1I). Meanwhile, the helix consisting of residues methionine 1 through serine 34 is stabilized by loose non-native contacts with the DNA-binding region (Fig. S1H), as the C-terminal helices with which it pairs to form the dimerization region have not yet been synthesized. These helices have been partially synthesized by length 112, but dimerization-region folding is still unfavorable at this point. The entirety of the C-terminal helices must be synthesized, which occurs around the full monomer length of 144, for the dimerization region to acquire partial stability (≈ 70% folded at the temperature shown.) We note that these results are reported at a simulation of temperature of *T* = 0.51 *T*_*M*_, where *T*_*M*_ is the DNA-binding region melting temperature. We chose this temperature because it is slightly below the dimer melting temperature of *T* ≈ 0.65 *T*_*M*_ (Fig. 2B) and corresponds to a physiologically reasonable folding stability of ∼ 5 *k*_*B*_*T* (Fig. S1B). However, our results are consistent across temperature choices below the dimer melting temperature (Fig. S1E). We further note that, although real physiological temperatures typically lie only slightly below protein melting temperatures, our temperature choice of *T* = 0.51 *T*_*M*_ is nonetheless reasonable in our model because our potential energy function is temperature-independent.

### MarR folding rate rapidly decreases beyond 100 amino acids due to non-native interactions

We next asked how the folding kinetics for MarR’s rate-limiting folding step, namely DNA-binding region folding, change as the nascent chain elongates beginning at 100 amino acids. We find that for a narrow window around this length, the rate-limiting step is both thermodynamically favorable and relatively fast (Fig. 2D). Beyond 100 amino acids, this step becomes dramatically slower. By length 112, this rate has decreased by roughly 1000-fold, and by the time the monomer is fully synthesized (144 AAs), the rate has decreased by roughly 2000-fold relative to the 100 AA partial chain (Fig. 2D, bottom). This slowdown far exceeds what is predicted from general scaling laws of folding time as a function of length (1, 28, 29). For instance, the power law scaling proposed by Gutin et al. (29),*τ* ∼ *L*^4^, predicts only a ∼4-fold slowdown between lengths 100 and 144 AA. The discrepancy between this general scaling and our observed dramatic slowdown suggests that factors specific to MarR are at play. One possibility is non-native intermediates. To test this hypothesis, we turned off the contribution of non-native contacts to the potential energy by re-running simulations in an all-atom Go potential in which only native contacts contribute (30, 31). In stark contrast to the full knowledge-based potential (Fig. 3A, left), the native-only potential predicts that below the melting temperature, the full protein folds dramatically *faster* than the partial chain at length 100. Furthermore, whereas the full potential predicts that both folding rates drop with decreasing temperature, the native-only potential predicts that the folding rates remain constant or *increase* with decreasing temperature. These findings can be explained by two effects related to non-native contacts, namely 1.) The partial chain is normally stabilized by loose non-native contacts, and so their absence leads to a reduced thermodynamic driving force for folding (Figs S1H and S2E), and 2.) The absence of non-native contacts eliminates kinetic trapping for the full protein at low temperatures. As a result, the folding rate now increases, rather than decreases with lowering temperature due to a stronger thermodynamic driving force. These observations point to the importance of non-native interactions in producing the observed orders-of-magnitude slowdown in MarR folding rate in the full potential at lengths beyond 100 amino acids.

As an additional test of the role of non-native contacts, we examined snapshots that have yet to undergo the rate-limiting step and identified ones that are kinetically trapped, defined as having ≥ 5 non-native contacts that need to be broken before the rate-limiting step can occur. Snapshots that do not fulfill this criterion are deemed non-trapped, and generally take on a looser, more molten-globule like structure. We then computed the free energy difference between these trapped and non-trapped ensembles as a measure for the stability of misfolded kinetic traps (Fig. 3B). For all temperatures below the melting temperature, this free energy difference is greater for the MarR chain at length 100 than for the full protein. We note that at temperatures below *T* ≈ 0.85 *T*_*M*_, non-trapped structures are observed extremely infrequently, leading to large errors in this free energy calculation. We thus do not plot these temperatures. But the trend at temperatures above *T* ≈ 0.85 *T*_*M*_ clearly suggest that the full protein experiences deeper kinetic traps. Although we define trapped snapshots here as ones that have ≥ 5 non-native contacts, our results are robust to the choice of this threshold value (Fig. S2F).

**Fig. 3.**
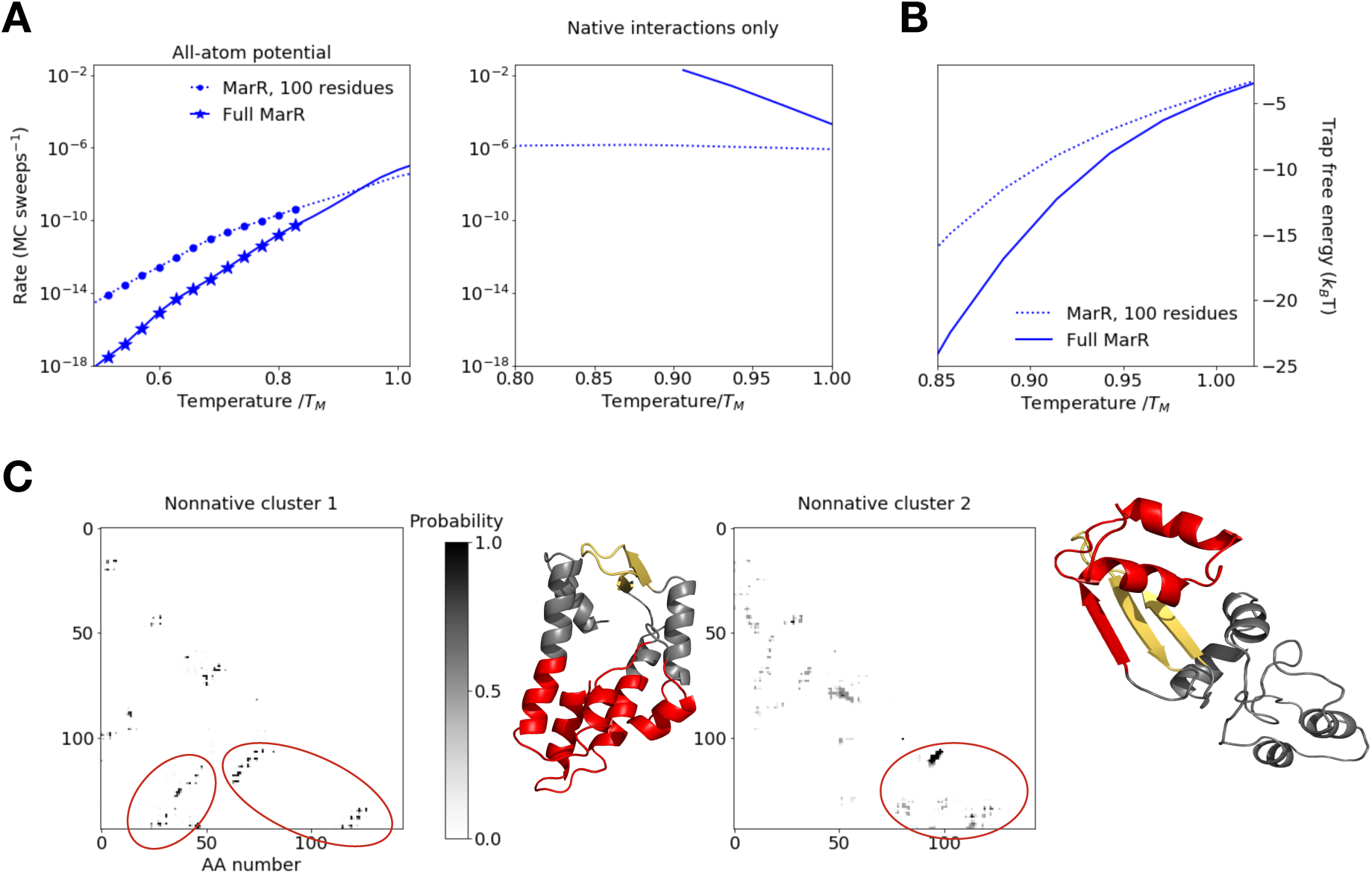
A) Folding rate vs temperature for DNA binding region folding rate as a function of temperature at nascent chain length 100 (dashed line) and full MarR (solid line), using the all-atom potential (left) and a native-central potential in which non-native interactions have been turned off (right). Symbols indicate temperatures at which the partial chain folds significantly faster than the full monomer (*p<* 0.01) based on bootstrapped distributions (see Methods) (B) Free-energy difference between configurations prior to the rate-limiting step that are kinetically trapped (defined as having at least 5 nonnative contacts that must be broken before rate-limiting step can occur) and those that are not trapped as a function of temperature for both the partial MarR chain at length 100 and full MarR. (C) Mean nonnative contact maps for the two most prevalent clusters (see Methods) among full MarR simulation snapshots in which the DNA binding region is not folded, along with representative structures. Contacts involving the C-terminus that most be broken before folding can proceed are circled in red on the maps and highlighted on the respective structures.

Since kinetic traps are deeper at chain lengths beyond 100 amino acids, we hypothesized that non-native contacts involving residues at sequence positions beyond 100 crucially stabilize these traps at longer lengths. To test this, we constructed and clustered the non-native contact maps of full protein snapshots prior to the rate-limiting step (see Methods), and visualized average non-native contact maps for these clusters (Fig. 3C). Indeed, the two most heavily populated clusters contain multiple non-native contacts involving amino acids beyond 100. In the first cluster (left), residues 51-55, which natively pair with the beta strand 95-100, are instead sequestered into a non-native hydrophobic core that is stabilized by C-terminal residues. In the second cluster (right), the beta strand 95-100 forms a non-native hairpin with residues 106-111, again impeding the native insertion of residues 51-55. Notably, many of the residues involved in stabilizing these non-native traps, particularly cluster 2, are already synthesized at length 112, thus explaining why the rate of folding is already much slower at that length than at length 100. Together, these contact maps further highlight the importance of C-terminal non-native contacts in drastically slowing folding as the nascent MarR chain elongates.

### Kinetic modeling predicts that vectorial synthesis helps MarR circumvent deep kinetic traps

Given that nascent MarR folding is fastest at chain lengths around 100 AAs, we hypothesized that vectorial synthesis may significantly improve folding efficiency as compared to what would be possible with unassisted post-translational folding. To test this, we developed a kinetic model of co-translational folding (Fig. 4A, details in Methods). Our model assumes that co-translational folding can be characterized by a fixed number of length regimes, namely chain length intervals for which the folding properties are nearly constant and informed by the calculations described above. For MarR, we identified three such regimes: 1.) 100-112 amino acids, at which point folding is relatively fast 2.) 112-144 amino acids, and 3.) 144 amino acids, corresponding to the full monomer. These latter two regimes both show similar folding properties, namely much slower folding and are depicted together as a single row in Fig. 4A. We assume that the protein spends a fixed amount of time at each length regime, during which it can fold or unfold as a continuous time Markov process (see Methods), prior to irreversible transition to the next regime via synthesis. This model contains two free parameters: 1.) The simulation temperature, which is kept at *T* = 0.51 *T*_*M*_ as before, and 2.) The ratio of the folding timescale to the synthesis timescale. This ratio cannot be determined from Monte Carlo simulations, which compute folding timescales in arbitrary Monte Carlo steps (although relative rates between different lengths or folding steps can be computed).

**Fig. 4.**
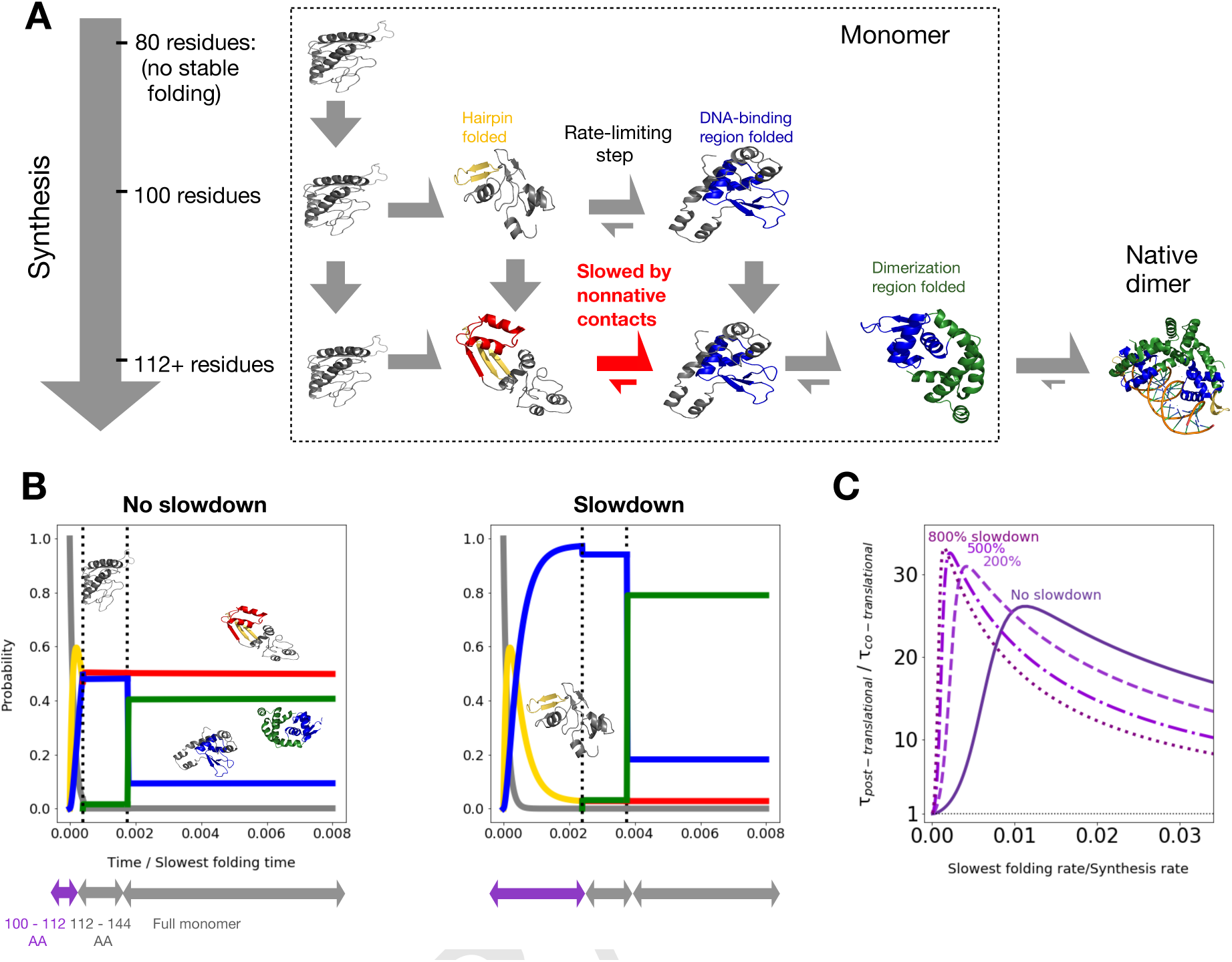
(A) Schematic of kinetic model (see main text and Methods for details). Dimerization is shown for completeness, but not accounted for in the kinetic model (B): Time evolution for the probability of occupying different states as a function of time, assuming the slowest folding rate is 6 · 10^−3^ times the protein synthesis rate (under constant translation speed). We further assume either no slowdown at conserved rare codons between residues 100-112 (left), or a 6-fold slowdown at rare codons (right, see main text and Methods). States are colored as in (A) (black = no native tertiary structure, gold = beta hairpin folded, red = beta hairpin folded with significant nonnative contacts, blue = DNA binding region folded, green = fully folded), and sample structures are shown. We neglect lengths prior to 100, at which point no folding occurs. (C) Fractional reduction in the mean time to complete synthesis and folding as a function of unknown synthesis rate, assuming various percent slowdowns at rare codons indicated by numbers over the curves and highlighted on the respective structures.

In Fig. 4b (left), we incorporate our computed folding rates for MarR into the kinetic model and plot the resulting probability of occupying different folding intermediates over time. We choose a set of parameters for which the effect of vectorial synthesis is particularly pronounced, namely we assume the slowest folding rate is 6 · 10^−3^ times the protein synthesis rate. For these parameters, enough time is spent at the 100-112 amino acid length regime that the DNA-binding region folds in roughly 50% of nascent chains (green and blue curves). The other half remains trapped in misfolded states (red curve). In contrast, an analogous simulation of post-translational folding shows no appreciable folding during this time period owing to the deep traps (Fig. S3A). Although vectorial synthesis is clearly advantageous, we wondered whether the advantage can be enhanced by slowing down MarR synthesis around the optimal folding length of 100. *In vivo* such a slowdown may result from a conserved stretch of rare codons which occurs roughly 30 amino acids downstream of this length (Fig. S3B). Indeed, we find that increasing the time spent in the 100-112 length regime by a factor of 6 increases the population that has undergone the rate-limiting step (green + blue curves) to nearly 100% (Fig. 4b, right). This suggests that, for these parameters, a rare-codon induced slowdown around length 100 significantly improves co-translational folding efficiency.

We next varied our model’s free parameters to test the generality of these results. In Fig. 4C, we show the mean time required for post-translational folding divided by the mean time for co-translational folding. This ratio is a proxy for the folding time benefit due to vectorial synthesis, with a value greater than 1 implying a benefit. We plot this ratio as a function of the unknown folding/synthesis timescale ratio, assuming that rare codons increase the time spent at the 100-112 length regime by various factors. We find that vectorial synthesis is always beneficial, although as expected this benefit diminishes as the folding/synthesis timescale ratio approaches zero, as the chain no longer has enough time to fold at length 100 (Fig. S3C). Furthermore, slowing down synthesis due to rare codons improves this benefit so long as the folding/synthesis timescale ratio is less than ∼ 0.01. For ratios above this, folding at intermediate lengths is fast enough that there is no benefit from slowing down synthesis (Fig. S3D). Thus in summary, our model predicts that 1.) for nearly all parameter values, MarR co-translational folding improves folding efficiency by helping nascent chains overcome deep kinetic traps, and 2,) assuming a reasonable range of timescales, rare codons tune synthesis rates so that a nascent MarR monomer can optimally exploit the faster folding rates available to it at lengths around 100 amino acids.

### Non-native interactions explain rare codon usage in multiple proteins

We then applied these methods to investigate the folding of other *E. Coli* proteins which were previously predicted to form stable folding intermediates upstream of conserved rare codon stretches (12). For each, we plot the native stability and the slowest folding rate as a function of chain length at a chosen temperature where the folding stability is physiologically reasonable (∼5-15 *k*_*B*_*T*). One example is the beta-ketoacyl-(acyl carrier protein) reductase, or FabG, an essential enzyme involved in fatty acid synthesis (Figs. 5A, S4). As with MarR, our simulations point to a rapid increase in monomer stability around 85 amino acids, at which point enough of the protein has been synthesized that a folding core composed of three N-terminal beta strands can fold (Fig. 5A top). This early folding step, which is rate-limiting overall, slows down somewhat beyond length 85, and even more beyond length 128, again owing to C-terminal non-native interactions (Figs. 5A bottom, S4 F-H). Thus, vectorial synthesis benefits FabG folding by allowing the chain to take advantage of these shorter lengths. The sequence contains various stretches of rare codons, each of which is predicted to potentially enhance this benefit under different conditions (Figs S4I-K). Another protein that shows similar behavior is the enzyme Cytidylate Kinase, or CMK (Figs 5B, S5). Our simulations predict that non-native kinetic traps lead to very slow CMK folding, consistent with previous experimental findings that the protein refolds on timescales of minutes (32). We further find that the stability notably increases with length at around 145 amino acids, even though our force field only predicts a folded fraction of ∼0.1 at this length. Slight inaccuracies in the force field may change this exact value, but our observation of a rapid increase in stability around this critical chain length is expected to be qualitatively robust. As with other proteins, this chain length corresponds to the point at which the rate-limiting step (beta-core nucleation) is fastest, as non-native contacts significantly slow the step at longer lengths (Figs 5B bottom, S5E-F). Furthermore, the chain-length window that corresponds to both increasing stability and relatively fast folding once again occurs roughly 30 amino acids downstream of a conserved stretch of rare codons (Fig. S5G). We note that, owing to large barriers in CMK’s landscape, the simulations did not converge adequately enough at low temperatures to allow for reliable folding rate calculations. We thus only compute folding rates at higher temperatures very close to the full protein’s melting temperature, at which point thermal stabilities are poor. However, we expect these trends to extend to lower, more physiologically reasonable temperatures, at which point the difference in folding rates, and thus the benefit due to vectorial synthesis, may be even more substantial.

**Fig. 5.**
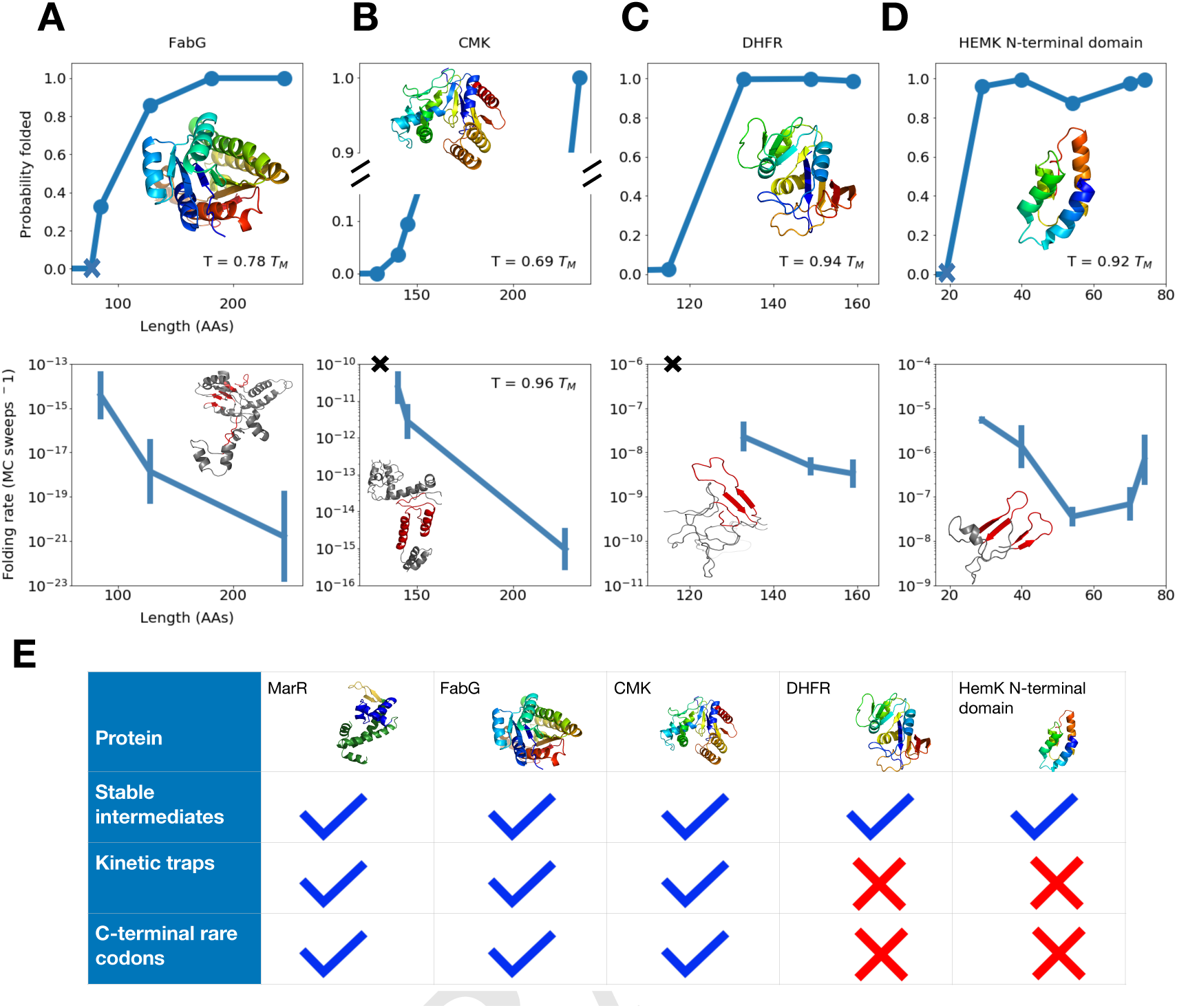
(A – D) As a function of chain length, the equilibrium probability that tertiary structure elements associated with the rate limiting step are formed (top) and the folding rate associated with the rate-limiting step (bottom) are shown for proteins (A) FabG, (B) CMK, (C) DHFR, and (D) HemK. For each protein, the native structure (top row) and a sample structure that has yet to undergo the rate-limiting folding step (bottom row) are shown, with C-terminal non-native contacts that must be broken prior to this step highlighted in red. Blue Xs’s in the top panels indicate the lengths at which the first amino acids associated with the rate-limiting step have been synthesized, while black X’s in bottom row indicate that no folding rate is computed because, even though enough residues have been synthesized for the rate-limiting structures to fold, their stability is low. As before, for each protein, we work at a temperature at which the fully synthesized chain shows a folding stability of ∼ 5 − 15 *k*_*B*_*T*. For more details pertaining to each protein, see SI. (E) For each protein simulated, we indicate if stable co-translational folding intermediates are formed, deep kinetic traps slow folding, and conserved C-terminal rare codons are found in the sequence.

### Counterexamples

Using our methodology, we also identified proteins for which vectorial synthesis and rare-codon induced pauses confer no benefit. We began by considering *E. Coli* Dihydrofolate Reductase (DHFR) (Figs. 3C, S6)—an essential enzyme which is known to fold rapidly (33–36). Indeed, our simulations predict no deep kinetic traps for full DHFR–the kinetic trap depth for unfolded states, computed as in Fig. 3B, is nearly zero at physiologically reasonable temperatures (Fig. S6F). Rather, the unfolded ensemble is characterized by loose, molten globule like states with significantly higher energy than the native state (Figs 3C bottom, S6E-G). Our predicted folding pathway (Fig. S6D) is in agreement with previous studies, which show that DHFR folds in multiple steps with fast relaxation times and no significant off-pathway intermediates (32, 33). Owing to this smooth folding landscape, we predict no advantage to vectorial synthesis, because even though the chain can fold at an intermediate length of 149, the folding kinetics hardly change with length (Fig. 5C). This is consistent with the protein’s codon usage: Although *E. Coli* DHFR contains C-terminal rare codons (Fig. S6H), they are not conserved and their synonymous substitution has been shown not to affect *in vivo* soluble protein levels nor *E. Coli* fitness (36). (However, conserved N-terminal rare codons were shown to be crucial for mRNA folding so as to ensure accessibility of the Shine-Dalgarno sequence (36).) In addition to DHFR, we simulated the N-terminal domain of HemK (residues 1-74, see Figs 5D, S7), a protein whose co-translational folding pathway has been studied using FRET by Holtkamp et al. (14). We find that the domain can adopt a stable native-like structure at around 40 amino acids, consistent with an observed increase in FRET near this length by Holtkamp and coworkers. But as with DHFR, slowing down synthesis at this length is predicted to confer no advantage (Fig. 5D), as the full domain folds rapidly and experiences only shallow folding traps at physiological temperatures (Fig. S7G). Consistent with this, the HemK N-terminal domain shows no conserved rare codons (Fig. S7H). Our results for every protein we simulate are summarized in Fig. 5b.

## Discussion

Together, these results shed light on how vectorial synthesis and its regulation affect the efficiency of i*n vivo* co-translational folding for various proteins depending on their nascent chain properties. The main takeaway is summarized in Fig. 6. For the relatively large single-domain proteins MarR, FabG, and CMK, we identify a narrow window of chain lengths at which folding is both favorable and fast. Prior to this length, the nascent chain cannot yet adopt native-like structures, while beyond this length, the folding rate drops by orders of magnitude. This dramatic drop in folding rate far exceeds what is expected due to increasing chain length alone (1, 28, 29) and instead results from deep non-native contacts involving C-terminal residues, which must be broken before folding can proceed. Thus, vectorial synthesis is predicted to significantly benefit folding, as it allows these proteins to exploit the narrow window of lengths at which the problematic C-terminal residues have not yet been synthesized and folding is fast. Under certain conditions, slowing synthesis at these critical lengths is necessary to give the chain enough time to fold, consistent with the presence of conserved C-terminal rare codons ∼30 amino acids downstream. In contrast to co-translational folding, post-translational folding is expected to be much less efficient for these proteins owing to misfolded states. Our results may also explain why other proteins lack conserved C-terminal rare codons. Namely for DHFR and the HemK N-terminal domain, we find that although co-translational folding is possible, it is not advantageous relative to post-translational folding because the full proteins fold rapidly without populating significant kinetic traps.

**Fig. 6.**
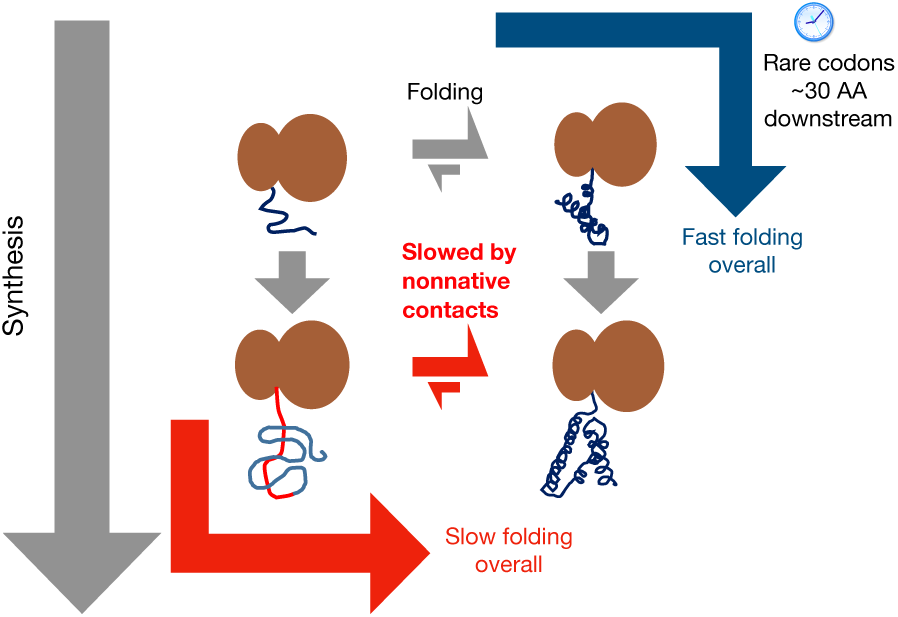
For misfolding-prone proteins that can fold co-translationally, the overall folding rate is optimized if the nascent chain has time to start folding at the earliest length at which stable folding can occur. At this point, the chain’s folding landscape is still relatively smooth (blue arrow). In case the nascent chain’s folding rate at this critical length is slightly slower than the synthesis rate, then slowing down synthesis using rare codons roughly 30 amino acids downstream is beneficial. In contrast, delaying folding until further synthesis is complete (red arrow) leads to deep kinetic traps stabilized by C-terminal residues, which significantly slow folding.

This study both generates specific experimental predictions, and also advances our general understanding of codon usage in proteins. For decades, it has been known that synonymous mutations which alter translation speed can affect the folding of large proteins, potentially reducing fitness (17) or exacerbating disease symptoms (37–39). However, the mechanism for these effects has not been established. Other studies have examined the role of evolutionarily conserved clusters of rare codons at domain boundaries, suggesting that these may give individual domains time to fold co-translationally (40). But more recent work has shown that conserved rare codons may be found at *any* chain length at which folding can begin, and not exclusively at domain boundaries (12, 13). These studies did not, however, establish a rationale for slowing down synthesis in the middle of a domain. Our work provides a potential mechanistic explanation for these observations, pointing to the crucial role of misfolded intermediates stabilized by C-terminal residues. In the cell, such intermediates may be involved in harmful aggregation, an effect that is not considered in our model but which may further heighten selection for co-translational folding. It is further worth noting that some rare codons, particularly at the 5’ end of genes, have evolved for reasons unrelated to co-translational folding, for instance to promote proper mRNA folding (36, 41, 42), or to minimize ribosome jamming (43). However, our work focuses on rare codons further downstream in coding sequences, at which point a nascent chain will be synthesized to a greater extent and co-translational folding becomes possible.

More generally, this work expands our understanding of how evolution optimizes the folding of large, misfolding-prone proteins *in vivo*. Besides vectorial synthesis and codon usage, another regulatory strategy involves chaperones. Growing evidence suggests that these two strategies may work in tandem in the cell, as chaperones such as trigger factor, DnaK, and TriC have been shown to bind nascent chains and promote co-translational folding (4, 8, 9, 44). Thus, rare codons may serve an additional role of slowing synthesis to give time for chaperones to bind. This may be especially beneficial if co-translational folding intermediates are non-native like, aggregation prone, or if these intermediates must undergo slow steps such as such as proline isomerization. Our method for studying co-translational folding, including the role of misfolded intermediates, can be applied in the future to shed light on these roles for chaperones, and potentially myriad additional factors that regulate protein folding *in vivo*.

## Materials and Methods

### Atomistic Monte Carlo simulations

Our algorithm for computing folding rates utilizes atomistic Monte-Carlo simulations with a knowledge-based potential and a realistic move-set comprising backbone and sidechain rotations (20–22). For each full protein construct and intermediate chain length, we performed the following steps:

1. A starting structure was downloaded from the PDB (PDB IDs for each protein shown in Table S1). This starting structure was equilibrated in the full potential for 15-30 million MC steps at a very low simulation temperature with harmonic umbrella biasing along native contacts. Umbrella biasing during equilibration increases the likelihood that the protein undergoes slight conformational changes relative to the starting structure that are necessary to attain the lowest energy configuration in the potential. Nascent chain constructs at intermediate lengths (for example, MarR at length 100) were then generated by truncating the C-terminus of the equilibrated full protein PDB structure, and equilibrating these truncated structures as was done for the respective complete protein.
2. To compute equilibrium thermodynamic properties, we ran replica exchange simulations using an added harmonic umbrella-sampling bias with respect to the number of native contacts. These simulations were run for 200-800 million MC steps at a wide range of temperatures. For some proteins, the initial 200-600 million MC steps additionally implemented a knowledge-based moveset (45) to aid the protein in finding energy minima at intermediate numbers of native contacts. However the timesteps that utilized these moves were not included in the free energy calculations, since these moves do not satisfy detailed balance.
3. To compute rates of unfolding, we ran simulations without replica exchange nor umbrella sampling at temperatures near or above the melting temperature. For all proteins, simulations were run starting from the equilibrated native structure. For FabG and CMK, we additionally ran unfolding simulations beginning from intermediate states containing a high degree of non-native structure, extracted from low temperature trajectories in the replica exchange simulations. Such simulations allow for a better estimate of the unfolding rate for these partially non-native intermediates at low temperatures.

### Simulation analysis and folding rate computation

To investigate a given construct’s folding properties, we first generated native contact maps of the respective fully synthesized and equilibrated structure, and identified islands of long-range contacts referred to as substructures (46). Native contact maps and substructures for each protein are shown in the SI. We then defined a coarse-grained folding landscape characterized by transitions between states defined by a subset of formed substructures. Such states are referred to as *topological configurations* (46). For fully synthesized MarR, example topological configurations include *abcdef* (all substructures folded), *abc* (only substructures a, b and c are folded) and Ø (no substructures folded–see Fig. S1). The resulting network of topological configurations is analogous to a Markov state model (47) in which states are defined based on structural features, rather than directly from kinetic information. This is justified because the folding/unfolding of a native substructure typically requires the forming/breaking of a loop, which is associated with a large free energy barrier. Thus, topological configurations show Markovian dwell-time distributions, as microstates consistent with a topological configuration rapidly equilibrate relative to the timescale of transition between topological configurations. (46).

Having defined substructures for a given protein, we assigned all simulation snapshots from replica exchange simulations to a topological configuration in accordance with which substructures are formed. Using the replica exchange simulations, we then used the MBAR method (48) to compute a potential of mean force (PMF) as a function of topological configuration–examples for MarR are shown in Fig. S1. The MBAR method was also used to compute PMFs as a function of number of native contacts or presence/absence of kinetic trapping (as in Fig. 3C) The PMF as a function of native contacts was used to compute a thermal average number of native contacts at each temperature, as in Fig. 2B.

To analyze unfolding simulations, we first assigned snapshots from these simulations to topological configurations, as above. To account for misclassification due to possible structural ambiguity, we fit the unfolding trajectories to a Hidden Markov Model that assumes a constant and uniform probability of misclassification to any incorrect configuration. We then identified clusters, or sets of topological configurations that are in rapid exchange. This was accomplished by defining a kinetic distance between topological configurations i and j, defined as the average time to transition between them, then clustering together configurations whose distance is below some threshold. The threshold was chosen to ensure a substantial separation between the timescales of exchange within the resulting clusters and exchange between clusters. This again ensures that clusters show Markovian dwell time distributions, which we have verified for MarR. The resulting clusters for each protein construct are shown in SI. Each snapshot from the unfolding simulations was then assigned to a cluster. At each unfolding simulation temperature, we then computed rates of unfolding between clusters, and fit the log rates as a function of temperature to the Arrhenius equation. Fig. S1 shows that the Arrhenius equation provides a good fit for the observed MarR unfolding rates. Using the Arrhenius equation, we then extrapolated unfolding rates to lower, more physiologically reasonable temperatures. We also computed the relative free energies of each cluster at those temperatures using the PMFs as a function of topological configuration obtained previously. From these unfolding rates and free energies, the folding rates between clusters were calculated from detailed balance. Namely, for two clusters i and j, the ratio of the forward and reverse transition rates *λ*_*i*→*j*_ and *λ*_*j*→*i*_ satisfies 

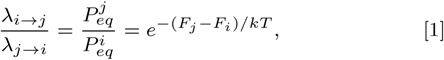

where *F*_*i,j*_ are the relative free energies of the respective clusters.

For each protein construct, we performed a bootstrap analysis to obtain an error distribution on folding rates by resampling 1000 times from the unfolding trajectories with replacement. We tested our method on HemK, for which folding transitions are fast enough for their rate to be directly calculated, and obtained good agreement (Fig. S7)

Using the PMFs as a function of topological configuration, we computed the equilibrium probabilities of forming structures associated with the rate-limiting folding step (Fig. 2D and Fig. 5) as follows: First, we identified the cluster that the protein transitions into during the rate limiting step. For MarR, this would be the cluster consisting of [*abc, bc, bcd*]. We then identified the substructures that are formed in the least folded configuration assigned to this cluster (*b* and *c* for MarR), and computed the Boltzmann probability that the protein occupies any configuration in which at least these substructures are formed. The minimum chain length at which the step can occur (colored Xs in these plots) was defined as the first length such that, for each of the substructures identified above, at least one native contact belonging to that substructure can form.

### Simulations with Native-only potential

These simulations for MarR at 100 residues and full MarR were run and analyzed as in the previous section, but with only native contacts found in the equilibrated structure contributing to the energy (30, 31). The values for attraction between native contacts, as well as added modest repulsion between non-native contacts, were tuned so that the ratio of the ground state energies of full MarR and MarR, 100 residues is close to that in the full knowledge-based potential.

### Clustering nonnative contact maps

To cluster misfolded states in accordance with which non-native contacts are present, we made nonnative contact maps of all snapshots assigned to a given topological configuration of interest at a set temperature range. The nonnative clusters for MarR in Fig. 3C include snapshots assigned to configuration *b*. We then assigned a distance between every pair of snapshots, defined as the Hamming distance between the contact maps (including only non-native contacts that are not present in the equilibrated native structure), and defined a distance threshold such that pairs of snapshots whose distance is less than this threshold are defined as adjacent. We formed clusters by finding the disconnected components of the resulting adjacency matrix. For most proteins, a distance threshold of 100 produced clusters that are structurally distinct and well-defined, but the results are robust to this precise value. Having defined clusters, we produced non-native contact maps for each cluster by averaging the contact maps of snapshots assigned to that cluster. Each resulting average contact map depicts the frequency with which non-native contact maps are observed in a given set of structurally similar misfolded states.

### Kinetic model of co-translational folding

To model co-translational folding, we defined a set of length regimes, each of which corresponds to an interval of chain lengths for which the protein’s folding properties are assumed to be constant. These folding properties are obtained by simulating a nascent chain at a length that is assumed to be representative of the length regime, and then applying the methods of the previous sections. At each length regime L, we define **P**^**L**,**T**^(*t*) as the vector of probabilities of occupying different clusters as a function of time at a given temperature T. Assuming continuous-time Markovian dynamics, **P**^**L**,**T**^(*t*) satisfies the master equation: 

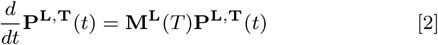

Where **M**^**L**^(*T*) is a transition matrix whose entries are given by 

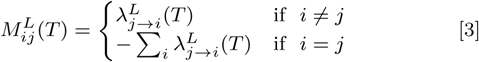

Where the folding/unfolding rates 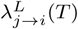 at length regime L are computed as described previously.

At each length L, the master equation is solved for an amount of time ·_*L*_ corresponding to the total time spent at length L, given an initial probability distribution **P**^**L**,**T**^(**0**). At the first length regime at which folding can occur, **P**^**L**,**T**^(**0**) is assumed to be one at the cluster containing the unfolded state (topological configuration Ø) and zero elsewhere. After time *τ*_*L*_, the probability **P**^**L**,**T**^(*τ*_**L**_) becomes the new initial distribution, **P**^**L**′,**T**^(**0**) at the next length regime *L*′, and the master equation is solved again given a new **M**^**L′**^ (*T*). In case cluster c at length L does not have an exact match at length L’, then for each cluster c’ at length L’, we define a similarity between c and c’ as the average number of substructures that must be formed or broken to transition from a topological configuration in c to one in c’. We then find the c’ that is most similar to c, and propagate element c of **P**^**L**,**T**^(*τ*_**L**_) to element c’ of **P** ^**L′**^,^**T**^ (**0**). The time spent at a given length regime *τ*_*L*_ is computed using: 

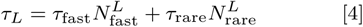

Where *τ*_fast_ and *τ*_rare_ are the average times to translate a fast and a rare codon, respectively, while 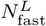 and 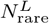 are the numbers of fast and rare codons in the length regime L. The values of *τ*_fast_ and *τ*_rare_ relative to characteristic folding times are unknown, and varied as free parameters as described in the main text.

In addition to computing how probability distributions evolve in time, we can compute the mean time to completion of synthesis and folding *τ*_total_ (Fig. 4C). To do this, we solve and propagate the probabilty distribution until the fully synthesized length regime F is reached, then evaluate the sum 

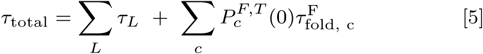

Where the second sum is over clusters in the full length 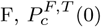 is the initial probability of occupying cluster c (obtained by propagating from the penultimate length regime as described above), and 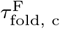 is the mean first-passage time to reach the cluster containing the folded cluster starting from cluster c. This mean first passage time is obtained by setting an absorbing boundary at the folded cluster and solving the equation: 

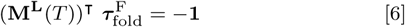

Where (**M**^**L**^(*T*))^T^ is the transpose of the transition matrix, 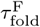 is a vector whose elements are the mean first passage times to the folded cluster from each initial cluster c, and the right hand side is a vector of negative ones.

## ACKNOWLEDGMENTS

The computations in this paper were run on the Odyssey cluster supported by the FAS Division of Science, Research Computing Group at Harvard University. AB was funded by the National Science Foundation GRFP (DGE1745303) and the Harvard Molecular Biophysics Training Grant (PI: James M Hogle, NIH/ NIGMS T32 GM008313). WMJ was funded by NIH grant F32GM116231. ES was funded by NIH grant R01 GM124044

## Supplementary Information

**Table S1:** List of PDB files used to simulate each protein

| Protein | PDB ID |
| --- | --- |
| MarR | 1JGS |
| FabG | 1Q7C |
| CMK | 2CMK |
| DHFR | 1DRA |
| HEMK | 1T43<br>Arginine 34 was replaced with Lysine to<br>match construct used in (1) |

**Figure S1:**
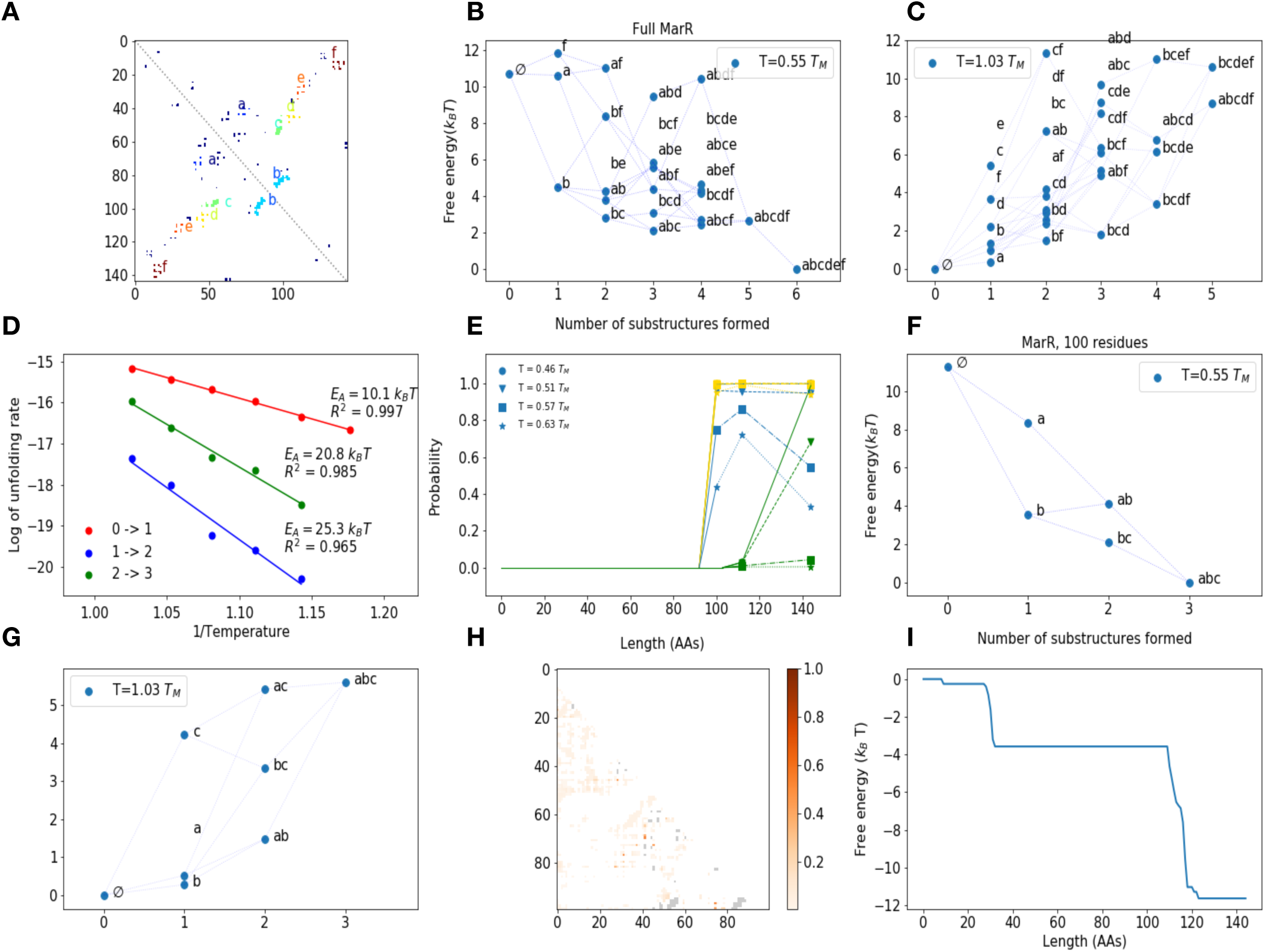
(A) Native contact map and substructures for MarR monomer. (B) and (C) Potentials of mean force (PMF) as a function of topological configuration for MarR at T = 0.55 T_M_ and T = 1.03 T_M_, where T_M_ is the DNA-binding region melting temperature. As the melting transition is crossed, configurations with less native structure become more favorable. (D) Sample Arrhenius plots for MarR showing that rates of transition between clusters, indicated in table S2. (E) Probability of forming minimal set of substructures associated with each folding step as a function of length as in main text Fig. 2D, at various temperatures. Colors are the same as in Fig. 2D, but different marker styles indicate different temperatures. As the temperature approaches the dimer melting temperature T = 0.65 T_M_, DNA binding region (substructures *b* and *c*) and dimerization region folding (substructures *a-d)* become less favorable, while the beta hairpin (substructure *b)* remains folded with high probability. But at all temperatures, a significant increase in DNA binding region stability is observed at length 100. (F and G) Same as (B) and (C) for 100 residue MarR nascent chain. The maximum substructures that can form at this chain length are *a, b*, and *c*. As shown in (F), the nascent chain at length 100 adopts a stable native-like topology (*abc*) at low temperatures. (H) Average nonnative contact map for snapshots of MarR, 100 residues assigned to topological configuration *abc*. The probability of each nonnative contact is indicated by color. Native contacts are shown in light gray in the background. (I) Minimum free energy relative to fully unfolded state as a function of chain length using the coarse-grained model in (2). A decrease in free energy around length 110 is observed that is analogous to our predicted rise in stability around length 100.

**Figure S2:**
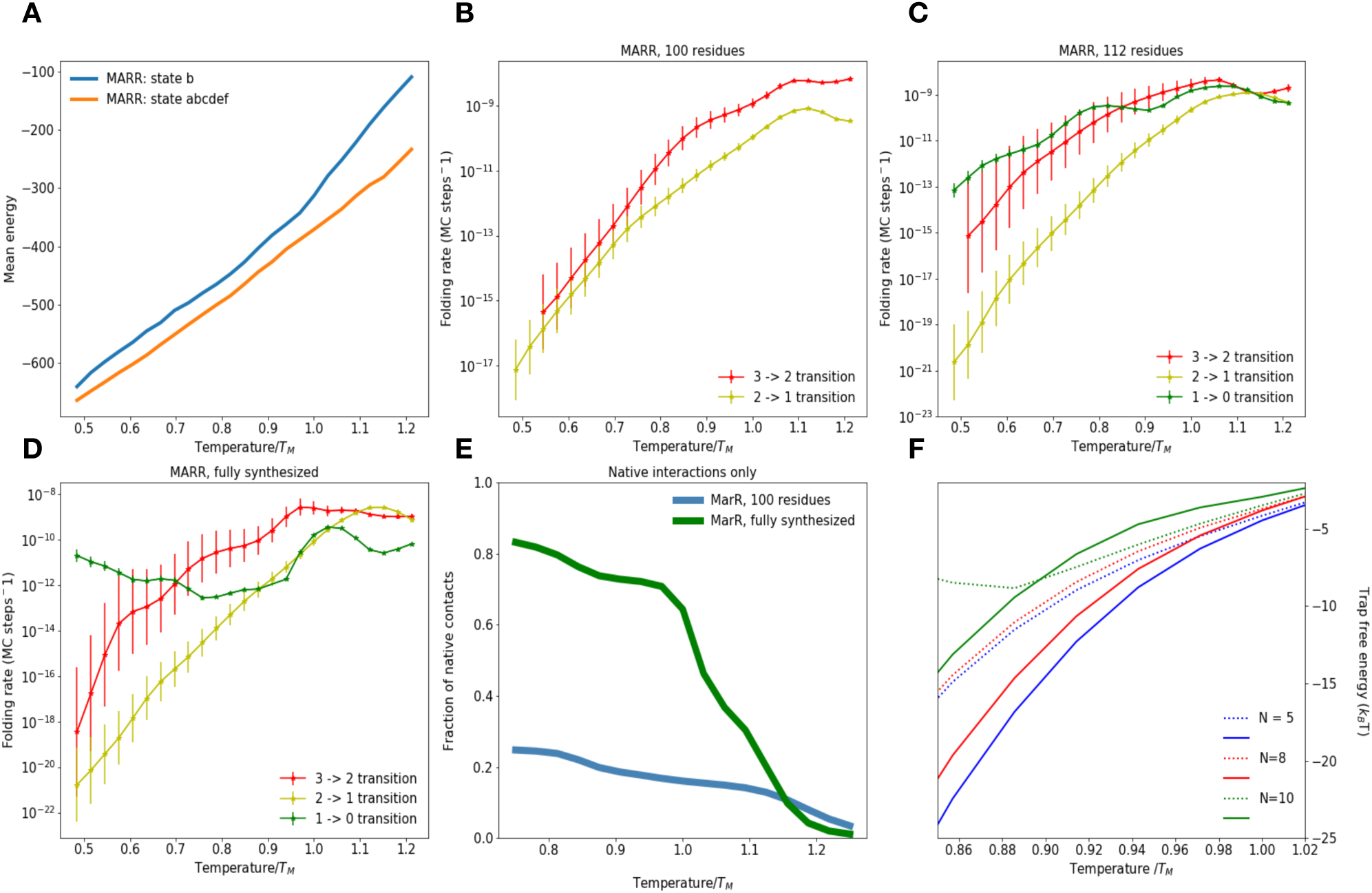
(A) Average energies of MarR snapshots assigned to topological configurations *b* (prior to rate-limiting step) and *abcdef* (maximally folded). A relatively small energy gap at low temperatures is indicative of non-native contacts stabilizing the b state. (B - D) Folding rates as a function of temperature for nascent MarR at chain length 100 (B), chain length 112 (C) and fuly synthesized monomer (D). In each panel, each line refers to a transition between a given pair of clusters (see methods). Topological configurations included in each cluster are listed in Table S2. For each transition, we only plot rates at temperatures for which the free energy difference between the clusters involved in the transition is less than 10 kT—for differences higher than this, statistical convergence of PMFs becomes poor. Error bars are obtained by bootstrapping (see Methods). (E) Fraction of native contacts as a function of temperature for MarR chain at length 100 and fully synthesized MarR as a function of temperature in the natives-only potential. The 100 residue chain shows worse stability than in the complete potential, where it is stabilized by non-native contacts. (F) Same as Fig. 3B, but for different values of N, the threshold number of non-native contacts that must be broken during rate-limiting step for a snapshot to be declared trapped (see methods). As in Fig. 3B, dashed lines represent MarR chain at length 100 while solid lines are full MarR. Each color represents a different threshold. For all thresholds, the full protein experiences deeper traps at temperatures below *T* ≈ 0.88 *T*_*M*_, indicating that this result is robust to the choice of threshold over a range of values.

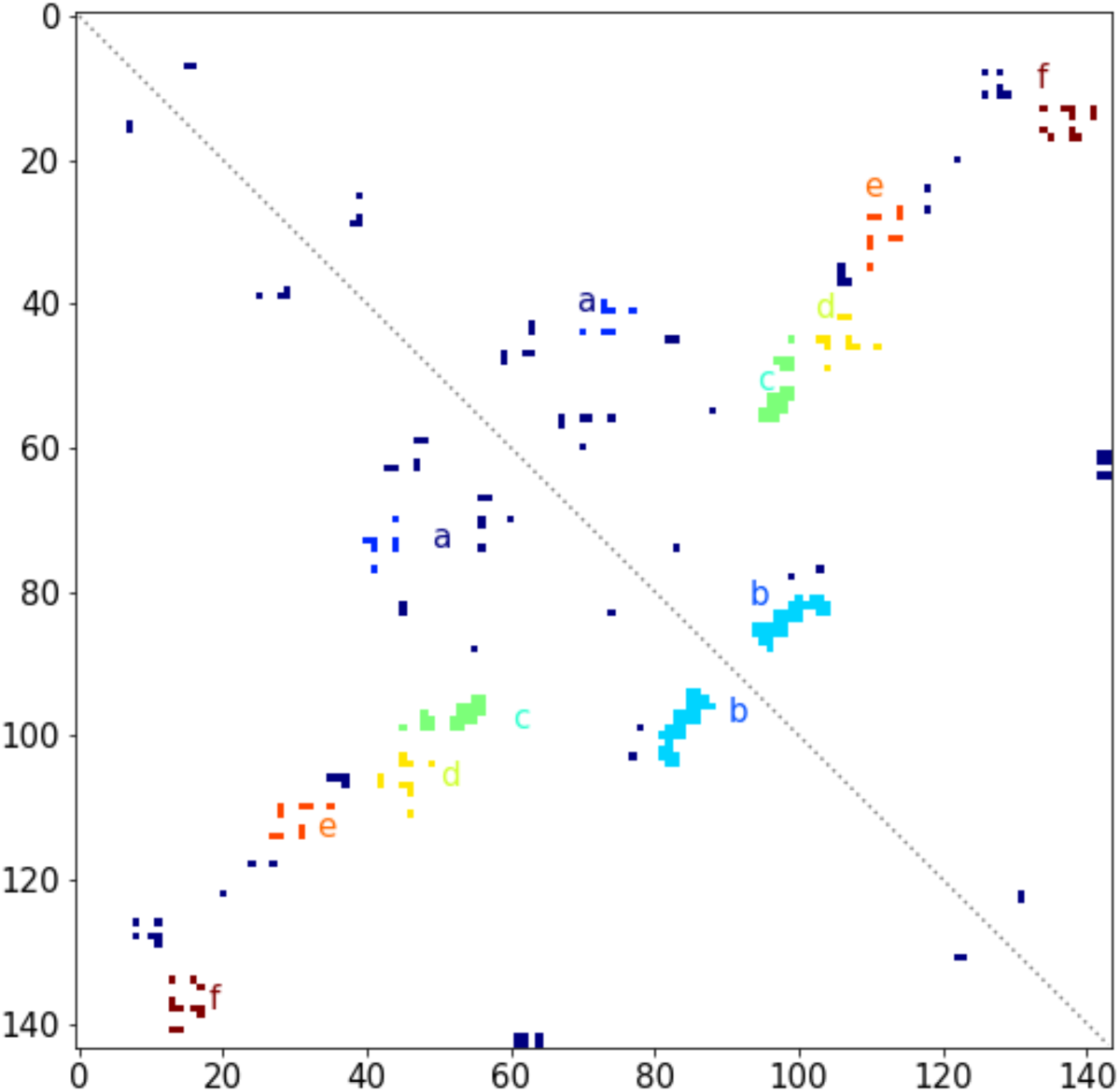

**Table S2:** Clusters for each MarR construct. Each cluster is defined as a set of topological configurations (listed above) that exchange quickly with one another relative to the timescale of exchange between clusters (see Methods). Native contact maps and substructures for MarR are shown above for reference. Other clusters that are not listed here are observed infrequently during unfolding simulations—these are not used for unfolding/folding rate calculations. For the full protein, we indicate which clusters are referred to in the text as having the beta hairpin region folded, DNA binding region folded, or being fully folded.

| Protein construct | Clusters |
| --- | --- |
| MarR, 100 residues | Cluster 1: [*abc, bc*]<br>Cluster 2: [*b*]<br>Cluster 3: [∅] |
| MarR, 112 residues | Cluster 0: [*abcde*]<br>Cluster 1: [*bcd, abc, bc*]<br>Cluster 2: [*b*]<br>Cluster 3: [∅] |
| MarR, fully synthesized (144 residues) | Cluster 0: [*abcdef, abcde, abcd*] (Fully folded)<br>Cluster 1: [*bcd, abc, bc*] (DNA binding region folded)<br>Cluster 2: [*b*] (Beta hairpin folded)<br>Cluster 3: [∅] |

**Figure S3:**
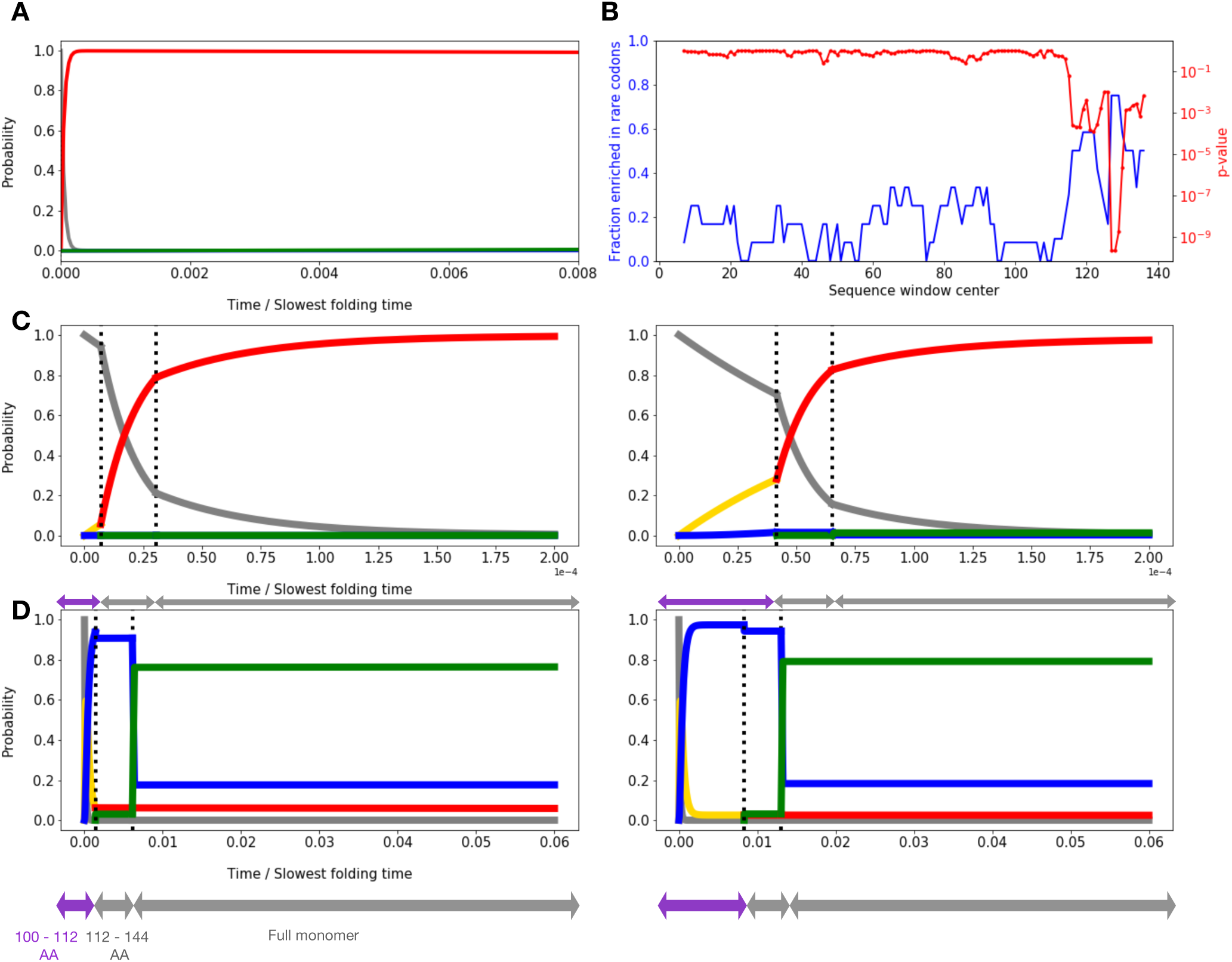
(A) Probability of occupying various MarR folding intermediates as a function of time assuming post-translational folding at T = 0.55 T_M_, for the same parameters and time period as in Fig. 4B. During this time period, nearly the entirety of the population remains kinetically trapped in the misfolded cluster 2 (red state with hairpin folded, but DNA binding region not folded). Color scheme is the same as in Fig. 4. (B) Fraction of homologous MarR sequences from sequence alignment enriched in rare codons as a function of sliding sequence window position, and associated p-value. Beginning around position 120, a large fraction of sequences contain rare codons. For details, see (2).(C) Same as main text Fig. 4B, except now assuming the slowest folding rate is 10^-4^ times the protein synthesis rate. Under this condition, folding is so slow compared to synthesis that the chain has insufficient time to fold co-translationally, even if rare codons are used. (D) Same as main text Fig. 4B, except now assuming the slowest folding rate is 0.02 times the protein synthesis rate (note change in x scale). Now, folding is fast enough that the protein folds co-translationally regardless of whether rare codons are used, so there is no benefit to slowing down. Arrows under plot indicate time spent in each length regime.

**Figure S4:**
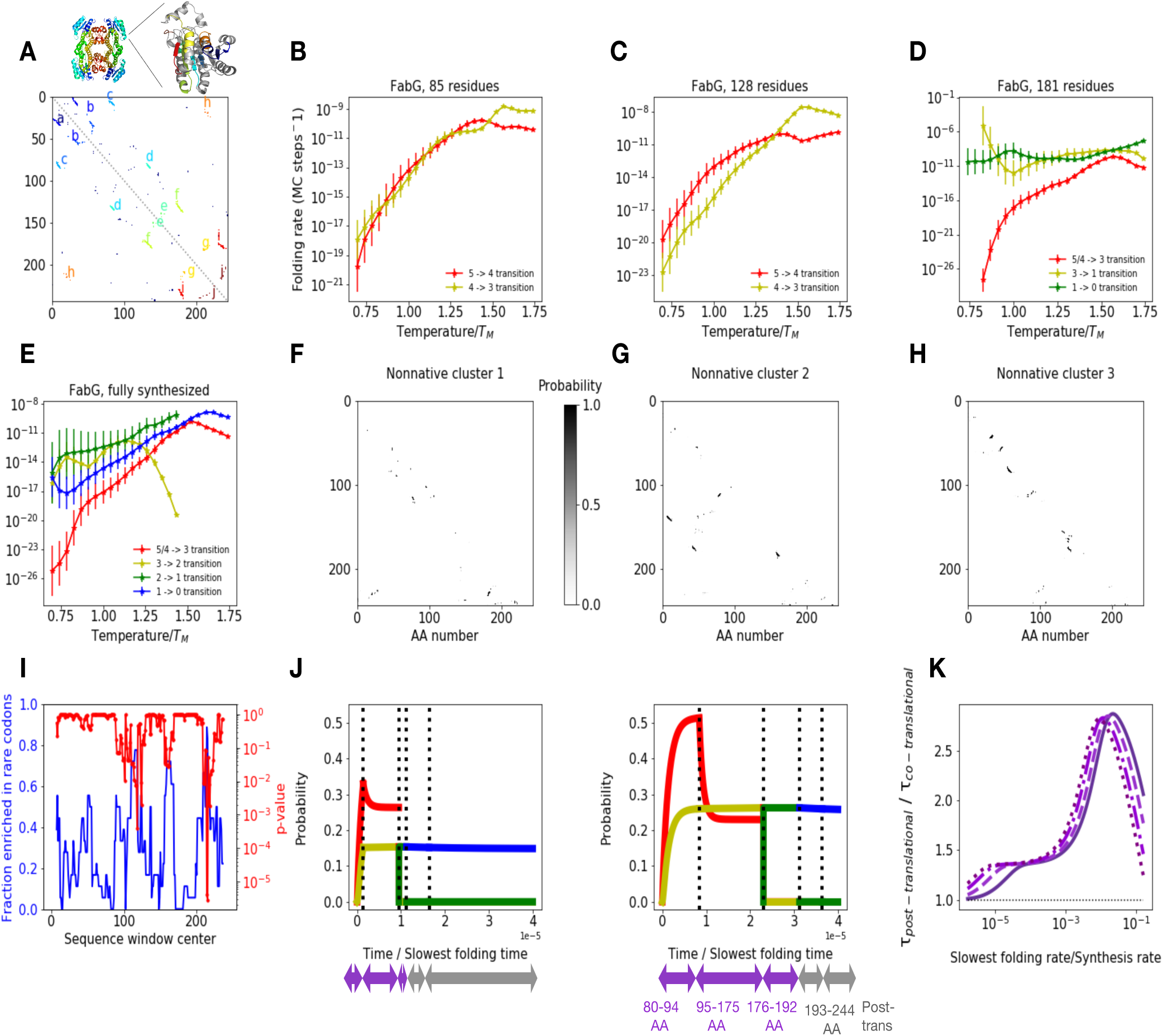
Summary of results for FABG (A) Native contact map and substructures for monomeric FABG. Crystal structures of the native tetramer and individual monomer are shown above the contact map. (B-E) Computed folding rate as a function of temperature at various nascent chain lengths for each transition. Topological configurations included in each cluster are listed in table S3. (F-H) Mean contact maps for the three most prevalent clusters among snapshots assigned to topological configuration A, prior to rate-limiting step. As with MarR, all clusters contain non-native contacts involving the C-terminus which must be broken before folding can proceed. (I) Fraction of homologous FabG sequences from sequence alignment enriched in rare codons as a function of sliding sequence window position, and associated p-value. In kinetic modeling, when rare codons are included, we introduce a slowdown in synthesis between AAs 80-94, 125-138, and 179-192 (roughly 30 amino acids upstream of each rare stretch). (J) Sample kinetic model results for probability of occupying various FabG folding intermediates as a function of time, assuming total protein synthesis time is ∼10^5^ times faster than slowest folding time and no slowdown at rare codons (left) and slowdown by factor of 6 at rare codons (middle). We consider the following length regimes (indicated under x axis): 80-94 AAs (assumed to have folding properties of 85 AA chain), 95-175 AAs (folding properties of 128 AA chain), 175-192 AAs (folding properties of 181 AA chain), 192-244 AAs, and post-translation (the latter two regimes have properties of full 244 AA protein). At each length regime, each curve corresponds to the population that has undergone the respective folding step shown in panels (E-H) from which the folding properties are derived indicated by the same color. (K) Reduction in mean first passage time to complete folding and synthesis relative to post-translational folding as a function of folding rate/synthesis rate ratio assuming various slowdowns at rare codons as in 4C (same colors). When folding is much slower than synthesis (ratio of ∼10^-6^ to 10^-4^), slowing synthesis is beneficial because the rare codon stretch centered around position 115 allows the chain to take advantage of fast folding at the 85 residue length regime. Note that the y values in this region are relatively low due to incomplete stability of the native-like intermediate at this length, which results in relatively low yield. For intermediate ratios between ∼10^-4^ and 10^-2^, the benefit due to co-translational folding increases, as the protein now has time to fold at the 128 amino acid length regime (where folding is slower than at length 85, but still faster than at full length). Slowing down synthesis is still useful, this time due to rare codon stretch centered around 155, which increases the time spent at the 128 amino acid length regime. For ratios of 10^-2^ and above, folding is fast enough that there is no need to slow down synthesis. Furthermore, the benefit due to co-translational folding starts to decrease due to this fast folding.

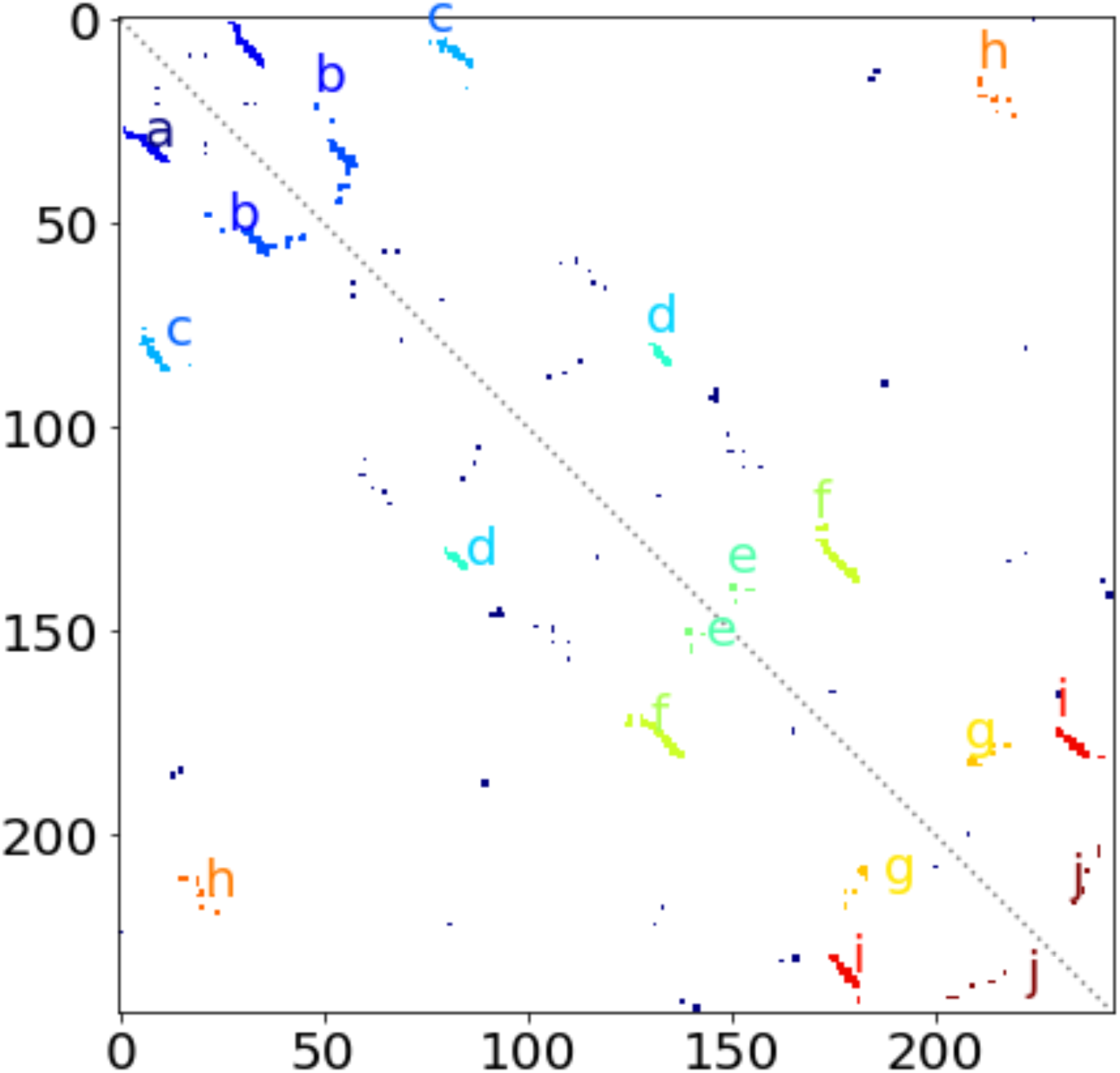

**Table S3:** Clusters for each FabG construct. In cases where the configurations assigned to a cluster at one chain length do not have an exact match at the subsequent length, we number clusters so as to indicate how population would be propagated to the next length based on structural similarity in kinetic model (see methods). For example, any population that occupies cluster 0 at length 181 are propagated to cluster 5 at length 244, even if the two clusters are not exactly alike. Likewise, any population in clusters 4 *or* 5 at length 128 are propagated to cluster 4/5 at length 181. These differences in cluster definition arise because at different lengths, different non-native contacts form during unfolding simulations, which dictate whether or not topological configurations are in fast exchange. We further note that for the fully synthesized FABG, the completely folded topological configuration is abcdefghij. However, we begin our unfolding simulations from state abcdef, since the fully folded state is thermodynamically disfavored when the protein is monomeric. We expect tetramerization will stabilize this fully folded state

| Protein construct | Clusters |
| --- | --- |
| FabG, 85 residues | Cluster 3: [ac]<br>Cluster 4: [a]<br>Cluster 5: [∅] |
| FabG, 128 residues | Cluster 3: [ac]<br>Cluster 4: [a]<br>Cluster 5: [∅] |
| FabG, 181 residues | Cluster 0: [abcdef, acdef, abcdf, acdf]<br>Cluster 1: [abcd, acd]<br>Cluster 3: [abc, ac]<br>Cluster 4/5: [∅] |
| FabG, fully synthesized (244 residues) | Cluster 0: [abcdef, acdef, abcdf, bcdf, acdf, cdf]<br>Cluster 1: [abcd, acd, cd]<br>Cluster 2: [abc]<br>Cluster 3: [ac]<br>Cluster 4/5: [a, ∅] |

**Supplementary figure 5:**
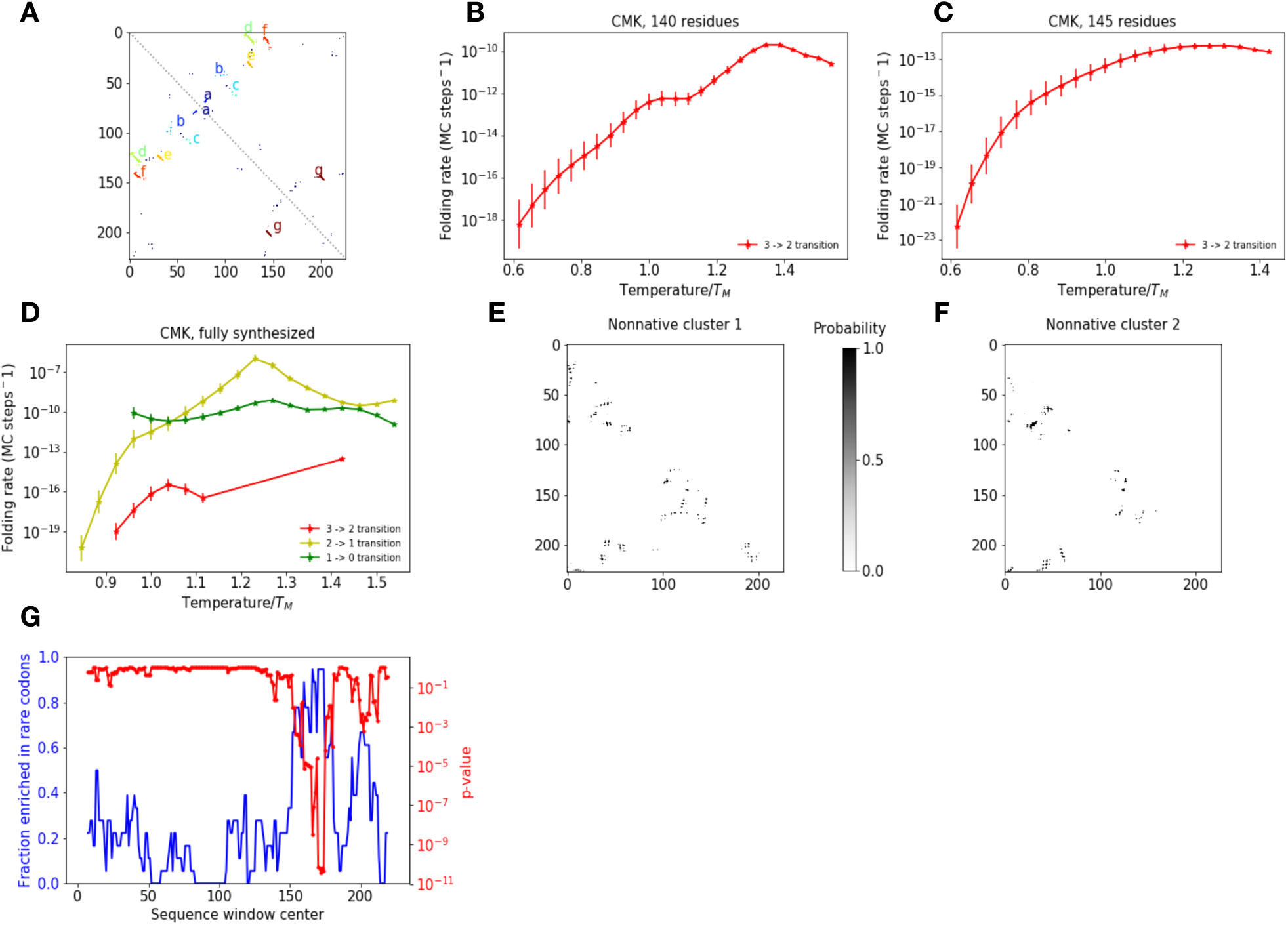
Summary of results for CMK: (A) Native contact map and substructures for CMK. (B-D) Computed folding rate as a function of temperature at various nascent chain lengths for each transition. Topological configurations included in each cluster are listed in table S4 (E-F) Mean contact maps for the two most prevalent clusters among snapshots assigned to topological configuration A, prior to rate-limiting step. As with MarR and FabG, both clusters contain non-native contacts involving the C-terminus which must be broken before folding can proceed. (G) Fraction of homologous CMK sequences from sequence alignment enriched in rare codons as a function of sliding sequence window position, and associated p-value

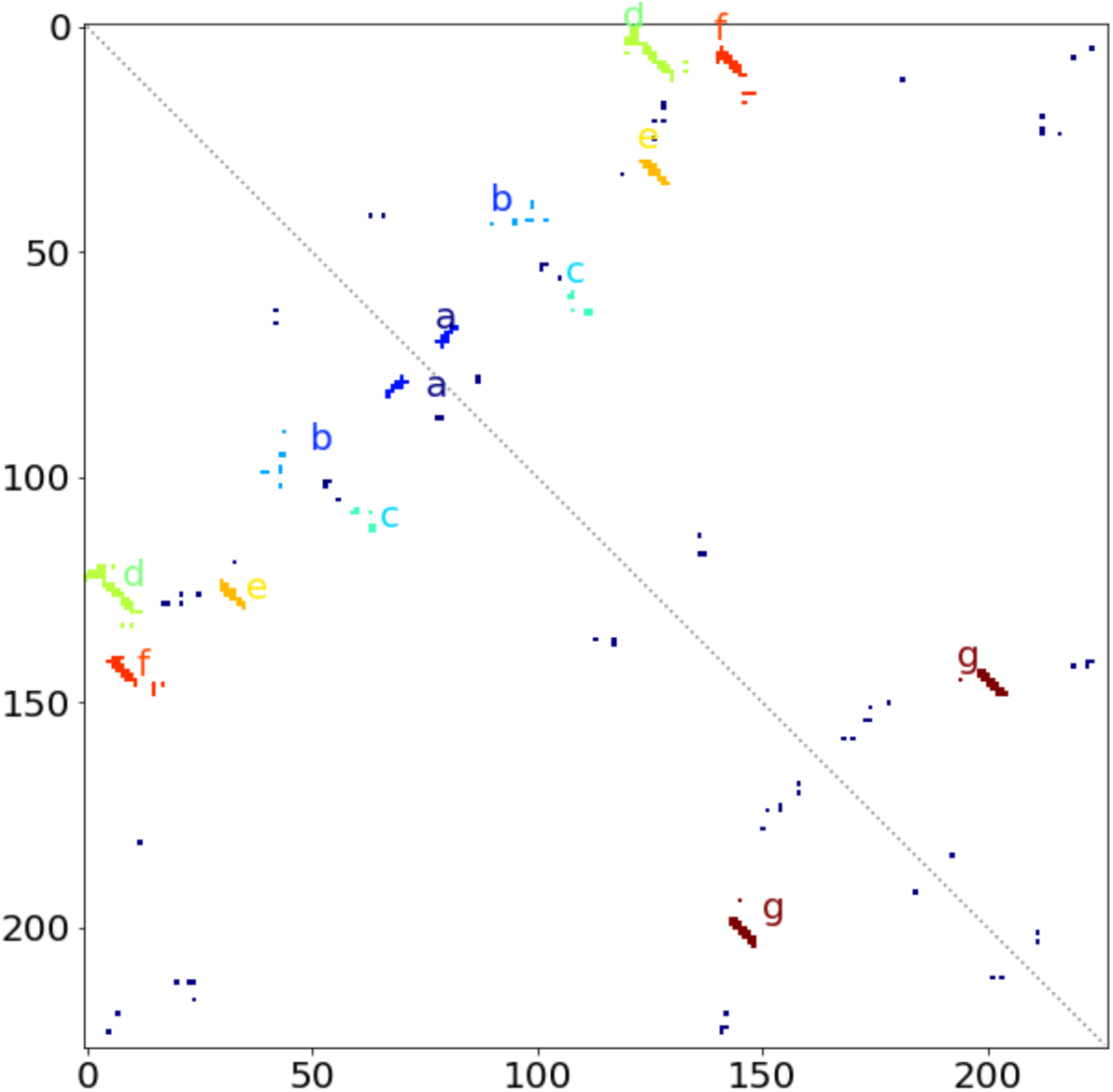

**Table S4:** Clusters for each CMK construct. We note that the first folding step involves the formation of substructure *a* (not computed), but this transition involves the simple folding of a short-range antiparallel beta hairpin and is not expected to be rate limiting. We further note that our PMFs predict that state acdefg is slightly lower in free energy at physiologically reasonable temperatures than the state abcdefg in which all substructures are formed, although these two differ by a relatively minor conformational change.

| Protein construct | Clusters |
| --- | --- |
| CMK, 140 residues | Cluster 2: [*abcde*, *acde*, *abcd*, *ade*, *ace*, *acd*, *ad*, *ac*]<br>Cluster 3: [*a*] |
| CMK, 145 residues | Cluster 2: [*acde*, *acde*, *ade*]<br>Cluster 3: [*a*] |
| CMK, fully synthesized (159 residues) | Cluster 0: [*acdefg*]<br>Cluster 1: [*acdef*, *adef*]<br>Cluster 2: [*ade*]<br>Cluster 3: [*ae*, *ad*, *a*] |

**Supplementary figure 6:**
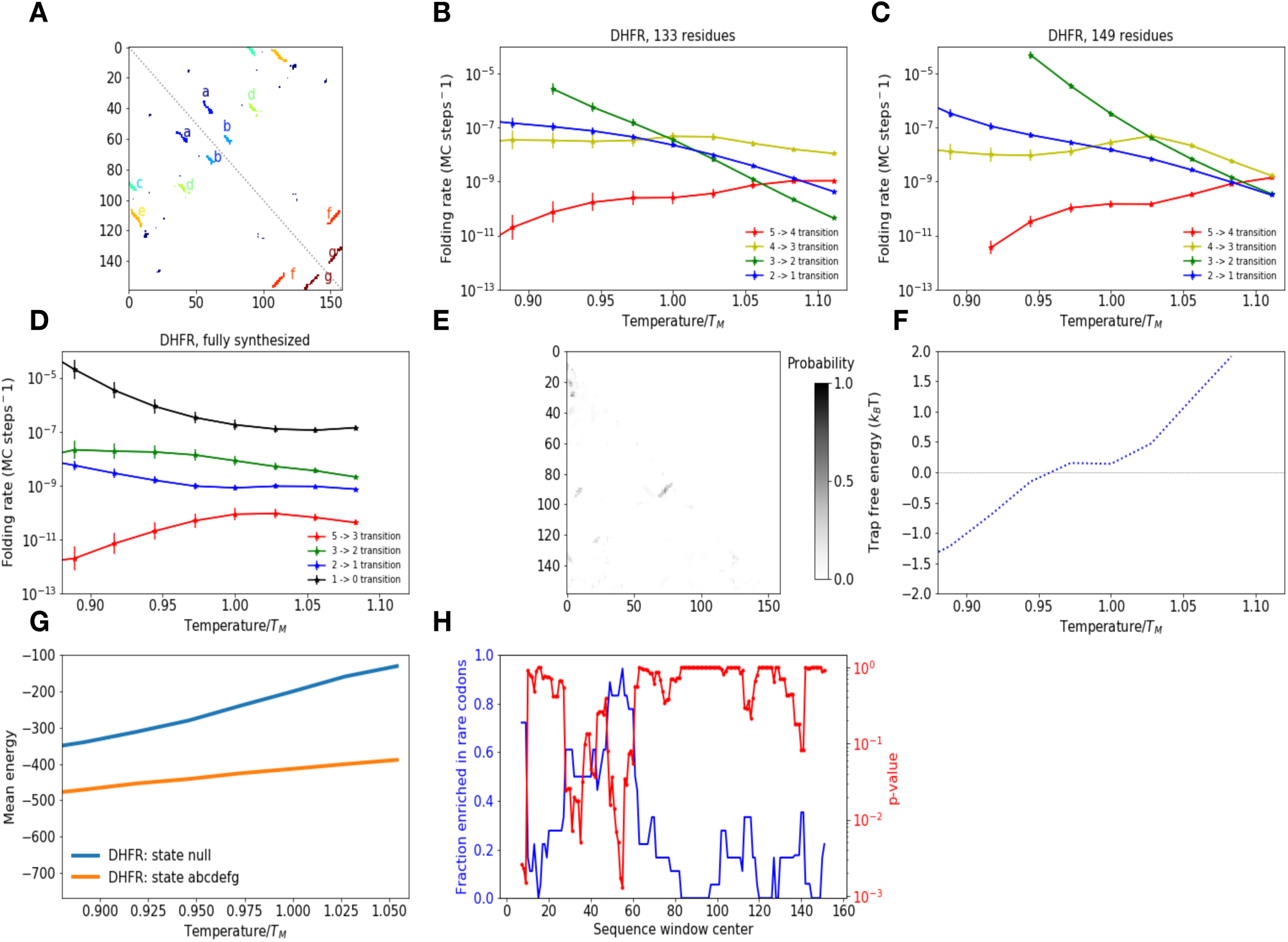
Summary of results for DHFR: (A) Native contact map and substructures for DHFR. (B-CD Computed folding rate as a function of temperature at various nascent chain lengths for each transition. Topological configurations included in each cluster are listed in table S5. (E) Mean nonnative contact map for snapshots assigned to Ø topological configuration (prior to rate-limiting step in fully synthesized DHFR). Nonnative snapshots cannot be readily clustered due to sparsity and lack of recurrence of non-native contacts. (F) Free energy difference between trapped and non-trapped subensembles that have yet to undergo the rate-limiting step in full DHFR (5->3 transition), defined as in main text Fig. 3b. At physiological temperatures around T = 0.9 T_M_, this free energy difference is nearly zero, indicating very shallow kinetic traps. (G) Average energy as a function of temperature for snapshots assigned to Ø and *abcdefg* (fully folded) states. The energy gap between these states is relatively large due to a lack of substantial non-native contacts. This is in contrast to MarR, where the energy gap is much smaller between states prior to the rate-limiting step and the folded state owing to substantial non-native contacts (Fig. S2A). (H) Fraction of homologous DHFR sequences from sequence alignment enriched in rare codons as a function of sliding sequence window position, and associated p-value. Although conserved rare codons are present at the N-terminus of the sequence, they are not found at the C-terminus.

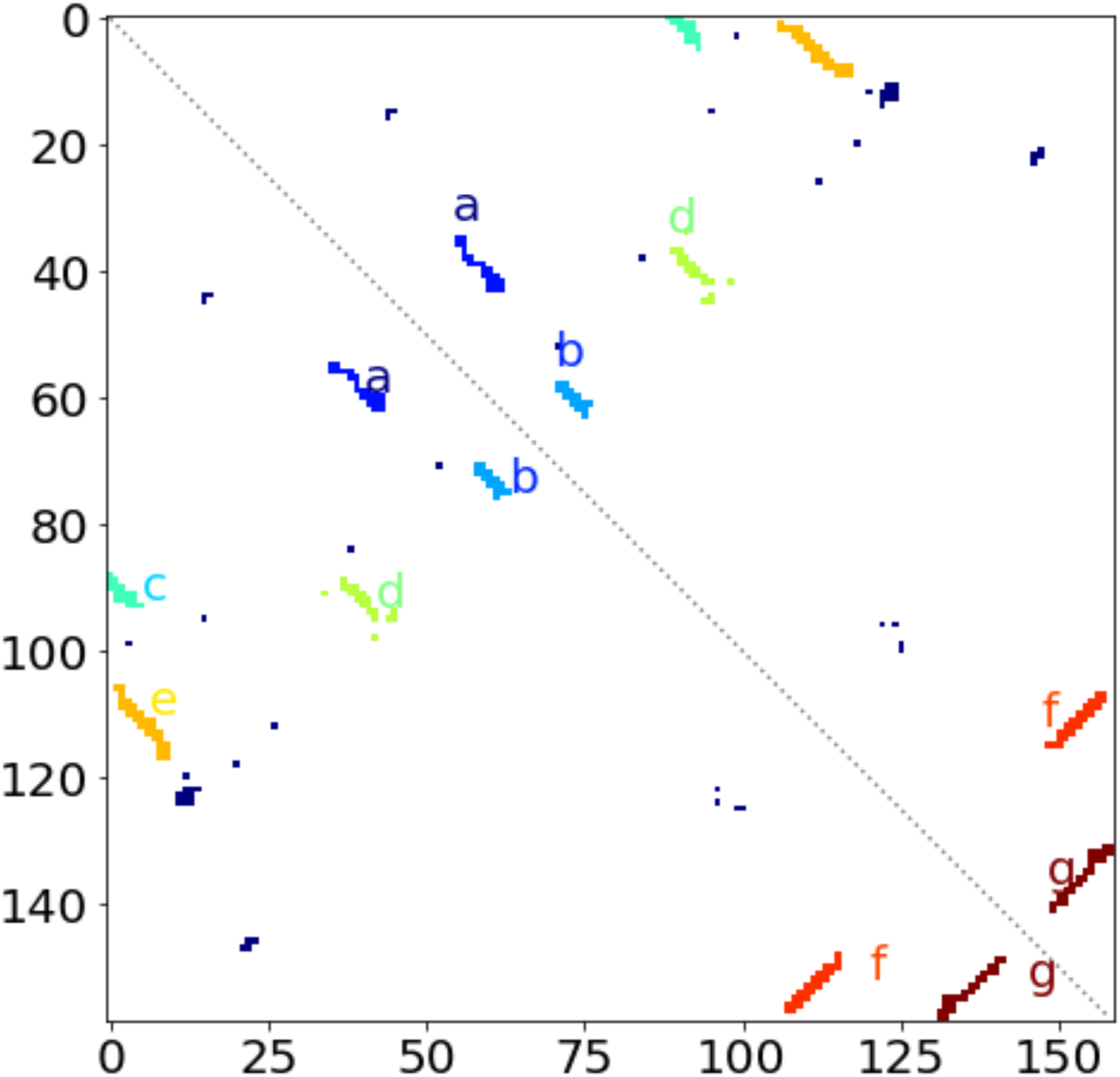

**Table S5:** Clusters for each DHFR construct. Note that although we did not construct a kinetic model for DHFR, if we did, cluster 4 at length 149 would be propagated to cluster 5 in the full protein.

| Protein construct | Clusters |
| --- | --- |
| DHFR, 133 residues | Cluster 1: $[abcde]$<br>Cluster 2: $[abcd]$<br>Cluster 3: $[abd, bd, ad]$<br>Cluster 4: $[ab, b]$<br>Cluster 5: $[\emptyset]$ |
| DHFR, 149 residues | Cluster 1: $[abcde]$<br>Cluster 2: $[abcd]$<br>Cluster 3: $[abd, bd, ad]$<br>Cluster 4: $[ab, b]$<br>Cluster 5: $[\emptyset]$ |
| DHFR, fully synthesized (159 residues) | Cluster 0: $[abcdefg]$<br>Cluster 1: $[abcde, acde]$,<br>Cluster 2: $[abcd, acd]$<br>Cluster 3: $[abd, ad, a]$<br>Cluster 5: $[\emptyset]$ |

**Supplementary figure 7:**
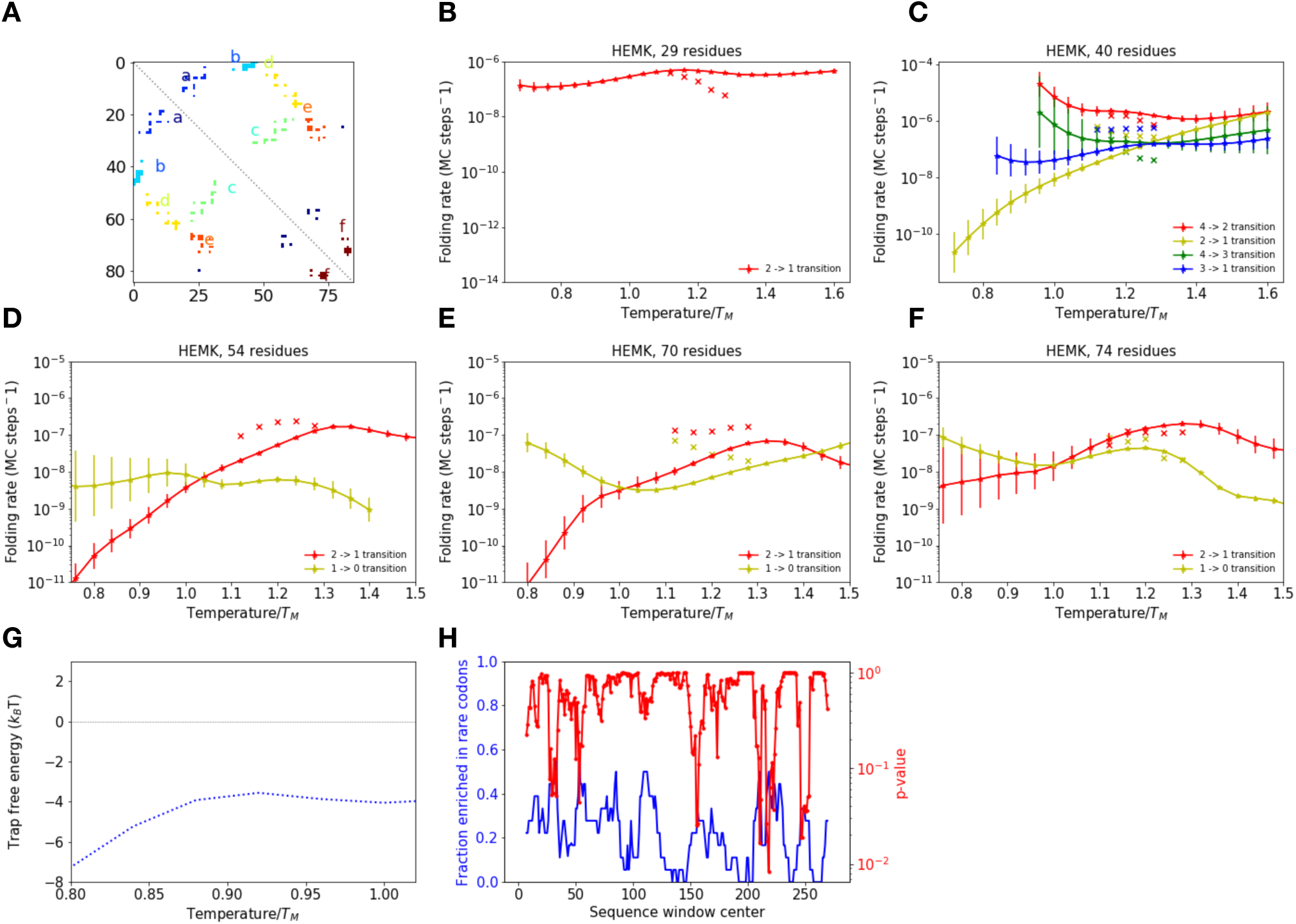
Summary of results for HemK N-terminal domain: (A) Native contact map and substructures for HemK residues 1-85 (however, we only simulate up to length 74). (B-F) Computed folding rate as a function of temperature at various nascent chain lengths for each transition. Topological configurations included in each cluster are listed in table S6. This protein is small enough that, for all these nascent chain lengths, our algorithm predicts that folding transitions are fast enough to be observable within a reasonable simulation timescale at the temperatures at which the unfolding simulations were run. Indeed, reversible unfolding/folding events are observed within the unfolding simulations. For each transition, we plot the observed refolding rates as Xs alongside the respective predicted rate. In most cases, the rates agree within an order of magnitude. Deviations typically result from either 1.) misclassification, whereby trajectories are falsely classified as having transiently refolded, or 2.) the presence of unfolding events that do not result misfolded states that are predicted to slow folding. At length 54, no 1->0 refolding events are observed, consistent with the predicted slow rate for this step. (G) Free energy difference between trapped and non-trapped subensembles that have yet to undergo the rate-limiting step at length 74 (2->1 transition), defined as in main text Fig. 3b. At physiological temperatures around T = 0.9 T_M_, this free energy difference is relatively small, around −4 k_B_T, as compared to the differences in excess of −15 k_B_T observed for MarR. This indicates relatively shallow traps for HEMK. (H) Fraction of homologous HemK sequences from sequence alignment enriched in rare codons as a function of sliding sequence window position, and associated p-value. No statistically significant conserved rare codons are found in the N-terminal domain (residues 1-74)

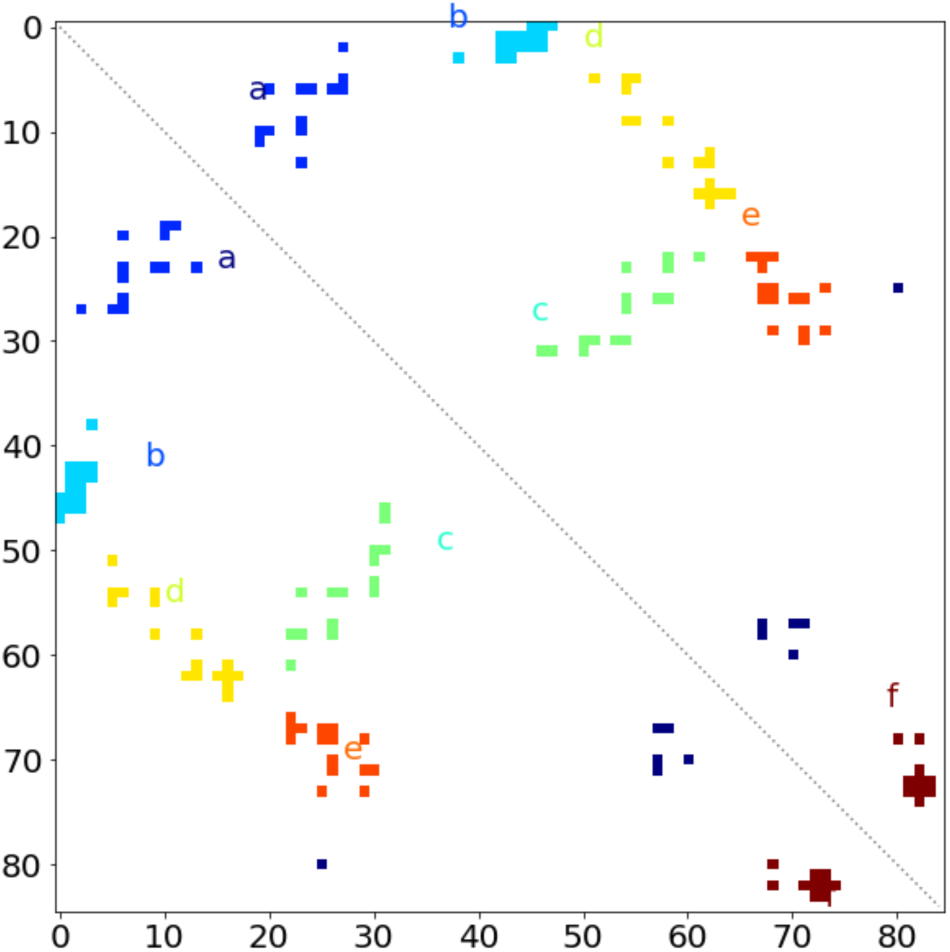

**Table S6:** Clusters for each CMK construct. We note that for lengths 29 and 40, we skip the clustering step based on kinetic connectivity in our analysis (see methods), as applying this step leads to clustering together of topological configurations that are unreasonably different in free energy at physiological temperatures. This is why more clusters are present at length 40 as compared to other lengths.

| Protein construct | Clusters |
| --- | --- |
| HEMK, 29 residues | Cluster 1: [a]<br>Cluster 2: [∅] |
| HEMK, 40 residues | Cluster 1: [ab]<br>Cluster 2: [a]<br>Cluster 3: [b]<br>Cluster 4: [∅] |
| HEMK, 54 residues | Cluster 0: [abcd]<br>Cluster 1: [abd, ab, a]<br>Cluster 2: [b,∅] |
| HEMK, 70 residues | Cluster 0: [abcde, abcd, abc, ab]<br>Cluster 1: [a]<br>Cluster 2: [∅] |
| HEMK, 74 residues | Cluster 0: [abcde, abcd, abc, ac]<br>Cluster 1: [abd, ab, a]<br>Cluster 2: [b,∅] |

